# Medicago SPX1 and SPX3 regulate phosphate homeostasis, mycorrhizal colonization and arbuscule degradation

**DOI:** 10.1101/2021.02.09.430431

**Authors:** Peng Wang, Roxane Snijders, Wouter Kohlen, Jieyu Liu, Ton Bisseling, Erik Limpens

**Affiliations:** Laboratory of Molecular Biology, Wageningen University & Research, 6708 PB Wageningen, the Netherlands

**Keywords:** SPX, phosphate homeostasis, arbuscular mycorrhiza, arbuscule development, strigolactone

## Abstract

To acquire sufficient mineral nutrients such as phosphate (Pi) from the soil, most plants engage in a symbiosis with arbuscular mycorrhizal (AM) fungi. Attracted by plant-secreted strigolactones, the fungi colonize the roots and form highly-branched hyphal structures called arbuscules inside inner cortex cells. It is essential that the host plant controls the different steps of this interaction to maintain its symbiotic nature. However, how plants sense the amount of Pi obtained from the fungus and how this determines the arbuscule lifetime is far from understood. Here, we show that *Medicago truncatula* SPX-domain containing proteins SPX1 and SPX3 regulate root phosphate starvation responses as well as fungal colonization and arbuscule degradation. *SPX1* and *SPX3* are induced upon phosphate starvation but become restricted to arbuscule-containing cells upon establishment of the symbiosis. Under Pi-limiting conditions they facilitate the expression of the strigolactone biosynthesis gene *DWARF27*, which correlates with increased fungal branching by root exudates and increased root colonization. Later, in the arbuscule-containing cells SPX1 and SPX3 redundantly control the timely degradation of arbuscules. This regulation does not seem to involve direct interactions with known transcriptional regulators of arbuscule degradation. We propose a model where SPX1 and SPX3 control arbuscule degeneration in a Pi-dependent manner via a yet-to-identify negative regulator.

## Introduction

In nature, plants usually face low mineral phosphate (Pi) availability in the soil, which limits their growth and development (Rouached et al., 2010). To deal with such phosphate limitation, plants typically induce a set of phosphate starvation induced (PSI) genes to acquire more Pi from the soil and to increase phosphate use efficiency (Bari et al., 2006; Zhou et al., 2008). In addition, most land plants engage in a symbiosis with arbuscular mycorrhizal (AM) fungi to increase their phosphate acquisition efficiency (Smith and Read, 2008). Under Pi starvation conditions, plant roots release strigolactones (SLs) that enhance spore germination and hyphal branching to initiate a symbiotic association (Akiyama et al., 2005; Besserer et al., 2006). Subsequently the fungus colonizes the roots and forms highly branched hyphal structures, called arbuscules, inside root cortex cells and its hyphae continue to form extensive networks in the soil. The hyphae can efficiently reach the scarcely available phosphate, which they deliver to the plant in return for carbon (fatty acids and sugars) (Luginbuehl and Oldroyd, 2017).

The maintenance of proper Pi homeostasis is important for plant growth and development, as either too low or too high Pi concentration in plant cells can be harmful to the plant (Wang et al., 2014). Therefore plants continuously sense and signal the phosphate status in response to their environment. Also during the symbiosis the plant must integrate phosphate status with fungal colonization and arbuscule development to keep the interaction beneficial. However, how a plant determines how much Pi it gets locally at the arbuscules and regulates Pi homeostasis in relation to arbuscule development is still an open question (Ezawa and Saito, 2018; Müller and Harrison, 2019).

Arbuscules form a symbiotic interface where Pi is provided to the host plant (Ezawa and Saito, 2018). They are relatively short lived structures that are degraded and removed from the cortical cells after 2-7 days (Kobae and Hata, 2010). This transient characteristic is thought to give the plant a means to locally abort arbuscules when these due to their age do not deliver sufficient nutrients (Lanfranco et al., 2018). In line with this, loss of the arbuscule-containing cell-specific PHOSPHATE TRANSPORTER 4 (PT4), responsible for transporting Pi across the peri-arbuscular membrane into the plant cell, leads to the premature degradation of the arbuscules in the model legume *Medicago truncatula* (Medicago) (Javot et al., 2007). This requires the activity of the MYB1 transcription factor, which induces the expression of hydrolytic enzymes such as cysteine proteases and chitinases to degrade the arbuscules (Floss et al., 2017). Intriguingly, it has been shown that Medicago can also adjust the amount of carbon that it delivers to the fungus depending on the amount of Pi obtained from the fungus (Kiers et al., 2011). AM fungal strains that deliver more Pi were shown to receive more carbon compared to less cooperative strains. This so-called reciprocal rewarding indicates that the plant is able to locally monitor the amount of Pi that it obtains from the fungus.

SPX domain containing proteins have emerged as key sensors and regulators of Pi homeostasis and signaling (Jung et al., 2018). The SPX domain is named after the Suppressor of Yeast gpa1 (Syg1), the yeast Phosphatase 81 (Pho81), and the human Xenotropic and Polytropic Retrovirus receptor 1 (Xpr1). This domain is able to sense the Pi status of a cell by binding with high affinity to inositol polyphosphates (PP-InsPs; (Wild et al., 2016; Jung et al., 2018). Changes in PP-InsPs levels in response to Pi deficiency are thought to modulate the activity of SPX-containing proteins and their interactors. The mode of action of single SPX domain containing proteins in the phosphate starvation response has been best studied in rice and Arabidopsis. The nuclear localized SPX1 and SPX2 proteins in Arabidopsis were shown to interact with PHOSPHATE STARVATION RESPONSE 1 (AtPHR1), a MYB transcription factor that together with its homologs controls phosphate starvation induced gene expression (Puga et al., 2014; Sun et al., 2016). Binding of AtSPX1/2 to AtPHR1 occurs at high Pi conditions and prevents the binding of AtPHR1 to the PHR1 binding site (P1BS) cis-regulatory element present in the promoters of many PSI genes. Similar regulation has been reported in rice where OsSPX1 and OsSPX2 control the activity of OsPHR2 in a phosphate dependent manner (Wang et al., 2014). Other SPX members, such as OsSPX4 have been localized in the cytoplasm where they control the cytoplasm-to-nucleus shuttling of OsPHR2 in a Pi dependent manner (Hu et al., 2019). Furthermore, OsSPX4 was reported to regulate nitrate and phosphate balance in rice through interaction with the nitrate transceptor OsNRT1.1B, which triggers OsSPX4 degradation upon nitrate perception (Hu et al., 2019). OsSPX3 and OsSPX5 have been localized in both the cytoplasm and nucleus, and redundantly modulate Pi homeostasis as functional repressors of OsPHR2 (Shi et al., 2014).

Given their key role in sensing and signaling of phosphate status in cells, we studied the role of SPX proteins during AM symbiosis. We identified two *SPX* genes that are strongly upregulated upon AM symbiosis, most specifically in the arbuscule-containing cells of Medicago. We show that these genes regulate Pi homeostasis in non-mycorrhizal conditions, and during the symbiosis positively control mycorrhizal colonization, in part through the regulation of strigolactone levels, as well as the timely degradation of arbuscules.

## Results

### SPX1 and SPX3 are strongly induced upon phosphate starvation and in arbuscule-containing cells

To identify *SPX* genes that might be important players during AM symbiosis, we first made a phylogenetic analysis of all single SPX domain proteins from Medicago, in relation to SPX proteins from arabidopsis and rice. This identified 6 members (Fig. 1A). We analyzed their expression in the roots of plants grown in high (500 μM) Pi, low (20 μM) Pi and 20 μM Pi plus *Rhizophagus irregularis* conditions for 3 weeks. qRT-PCR analyses showed that two *SPX* genes, *MtSPX1* (Medtr3g107393; hereafter *SPX1*) and *MtSPX3* (Medtr0262s0060; hereafter *SPX3*) were strongly induced by phosphate starvation as well as during the AM symbiosis (Fig. 1B). Transcriptome analyses of laser microdissected arbuscule-containing cells showed that these two SPX genes are dominantly expressed in arbuscule-containing cells (Fig. S1). To determine the spatial expression of *SPX1* and *SPX3* in more detail we analyzed promoter-GUS reporter constructs in transgenic roots grown for 3 weeks at 500 μM Pi, 20 μM Pi and 20 μM Pi with *R. irregularis* conditions. Plants grown at 500 μM Pi showed only weak GUS signal in the root tip, while under low Pi conditions strong GUS activity was detected throughout the root for both constructs (Fig. 1C-F). Sectioning of these roots showed that the SPX1 and SPX3 promoters were active in multiple cell types, including cortex and epidermis, of the root grown at low Pi, but not at high Pi (Fig. 1G, H). Upon AM symbiosis, the GUS signal became strongly restricted to the arbuscule-containing cells (Fig. 1I-L). No, or very low, GUS signal was observed in the non-colonized sections of the mycorrhizal roots. The arbuscule restricted expression was further confirmed by promoter:NLS-3×GFP analyses (Fig. S2). The same expression pattern of both *SPX* genes suggested that they may have similar functions, playing dual roles in the phosphate starvation response and AM symbiosis.

**Fig. 1.**
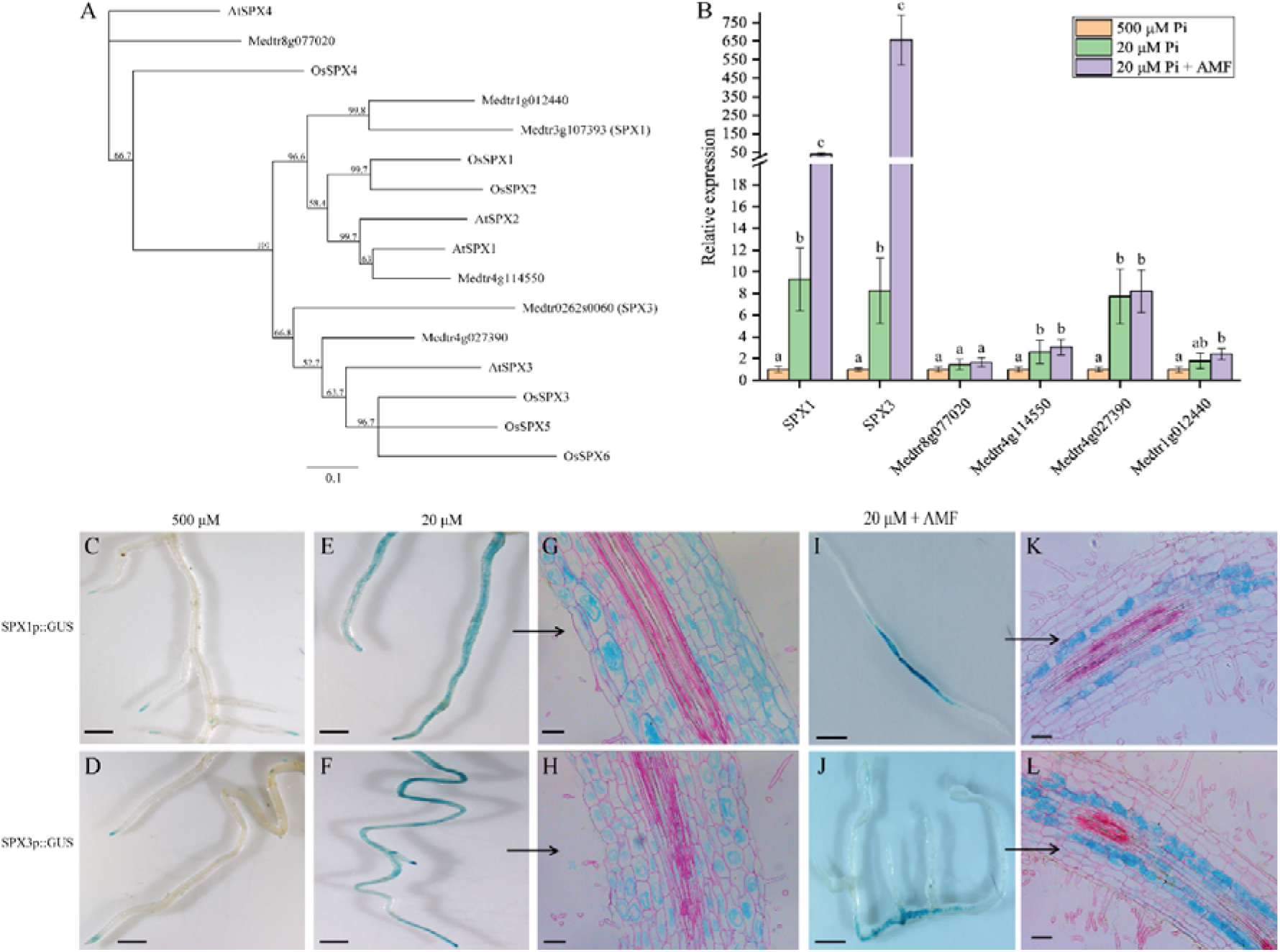
Medicago *SPX1* and *SPX3* induced by phosphate starvation and arbuscular mycorrhizal fungi. (A) Phylogenetic relationship of SPX proteins in Medicago, Arabidopsis and Rice. Unrooted tree constructed using Geneious R11.0 by neighbour-joining method with bootstrap probabilities based on 500 replicates. The identifiers of Arabidopsis and rice SPX proteins are listed in Table S4. (B) qRT-PCR analyses of Medicago *SPX* expression at high Pi (500 μM), low Pi (20 μM) and arbuscular mycorrhizal (20 μM Pi plus AM fungi (AMF)) conditions. *SPX1* (Medtr3g107393) and *SPX3* (Medtr0262s0060) are induced at low Pi conditions, and even stronger upon symbiosis with AMF. Medicago *Elongation factor 1* (*MtEF1*) was used as internal reference, error bars indicate standard error of the mean based on 3 individual plants. Different letters indicate significant differences between treatments (one-way ANOVA, P < 0.05). (C-D) *SPX1* and *SPX3* are expressed in root tips at 500 μM Pi. Scale bar =1 cm. (E-H) *SPX1* and *SPX3* at 20 μM Pi, (G) and (H) are longitudinal sections of (E) and (F). Sections are counterstained with 0.1% ruthenium red. Scale bar in (C-D), 1 cm, in (E-F), 100 μm. (I-L) *SPX1* and *SPX3* are highly and specifically induced in arbuscule-containing cortical cells (3 weeks post-inoculation). (K) and (L) are longitudinal sections of (I) and (J). Scale bar in (I-J), 1 cm, in (K-L), 100 μm.

To study the subcellular localization of SPX1 and SPX3, C-terminal GFP-fusion constructs were expressed in Medicago roots and Nicotiana leaves using either the constitutive *Lotus japonicus Ubiquitin1 (LjUB1)* promoter or their endogenous promoters. In all cases, including arbuscule-containing cells expressing SPX-GFP fusion under the control of their endogenous promoters, both fusion proteins localized to the cytoplasm as well as to the nucleus (Fig. S3A-E). The subcellular localization was not influenced by the Pi conditions (Fig. S3C, D).

To explain the transcriptional regulation of *SPX1* and *SPX3*, we examined their presumed promoter regions for known *cis*-regulatory elements involved in phosphate-starvation induced expression and arbuscule-specific regulation. This identified a P1BS (GNATATNC) binding site for the MYB transcription factor PHR (Bustos et al., 2010), a key regulator of PSI genes (Fig. S4). The presence of this element suggests that both SPX genes are under the control of PHR-dependent transcriptional regulation upon phosphate stress. Furthermore, we identified *cis*-regulatory AW-boxes (CG(N)_7_CNANG) and CTTC-elements (CTTCTTGTTC) in the promoter regions of both genes, which are binding sites for WRINKLED1-like transcription factors that have recently been found to regulate arbuscule-specific expression (Fig. S4). These elements can been found in the promoters of many arbuscule-enhanced genes, including genes involved in fatty acid synthesis and transport as well as genes required for phosphate uptake from the arbuscules (Xue et al., 2018; Jiang et al., 2018; Pimprikar et al., 2018). The presence of both P1BS and AW-box cis-regulatory elements may explain the observed expression patterns in the different conditions, although this remains to be experimentally verified.

### SPX1 and SPX3 regulate phosphate homeostasis

To study the function of SPX1 and SPX3, we identified *spx1* (NF13203_high_1) and *spx3* (NF4752_high_18) Tnt1-retrotransposon insertion mutants (Fig. S5A). Genotyping by PCR confirmed the Tnt1 insertion and a *spx1spx3* double mutant was developed by crossing *spx1* to *spx3* (Fig. S5B). RT-PCR confirmed the impairment of *SPX1* and/or *SPX3* expression in the respective mutant lines (Fig. S5C).

Since SPX proteins are thought to negatively regulate PHR activity at high Pi conditions to prevent the overaccumulation of phosphate, we first analysed the expression of PSI genes in the mutants and R108 wild-type under high and low Pi conditions. This showed a significantly higher expression of the PSI genes *Mt4* (U76742.1) (Burleigh and Harrison, 1999) and the phosphate transporter encoding gene *PT6* (Medtr1g069935) (Mbodj et al., 2018; Hu et al., 2019) in the *spx1spx3* double mutant under high Pi conditions (Fig. 2B). It correlated with a decreased shoot:root fresh weight ratio indicative of a phosphate starvation response in the double mutant (Fig. 2A, E). Furthermore, phosphate levels were increased in the shoots of the double mutant compared to wild-type plants grown at 500 μM Pi (Fig. 2F). Prolonged growth at 500 μM Pi further showed typical Pi toxicity effects (yellow colouring of the leaf margins) in the leaves of the double mutant (Fig. S6A). The single mutant lines did not show obvious phenotypes when grown at 500 μM Pi condition (Fig. 2A, E, F). These results suggest a negative role of SPX1 and SPX3 on phosphate starvation responses when ample Pi is available.

**Fig. 2.**
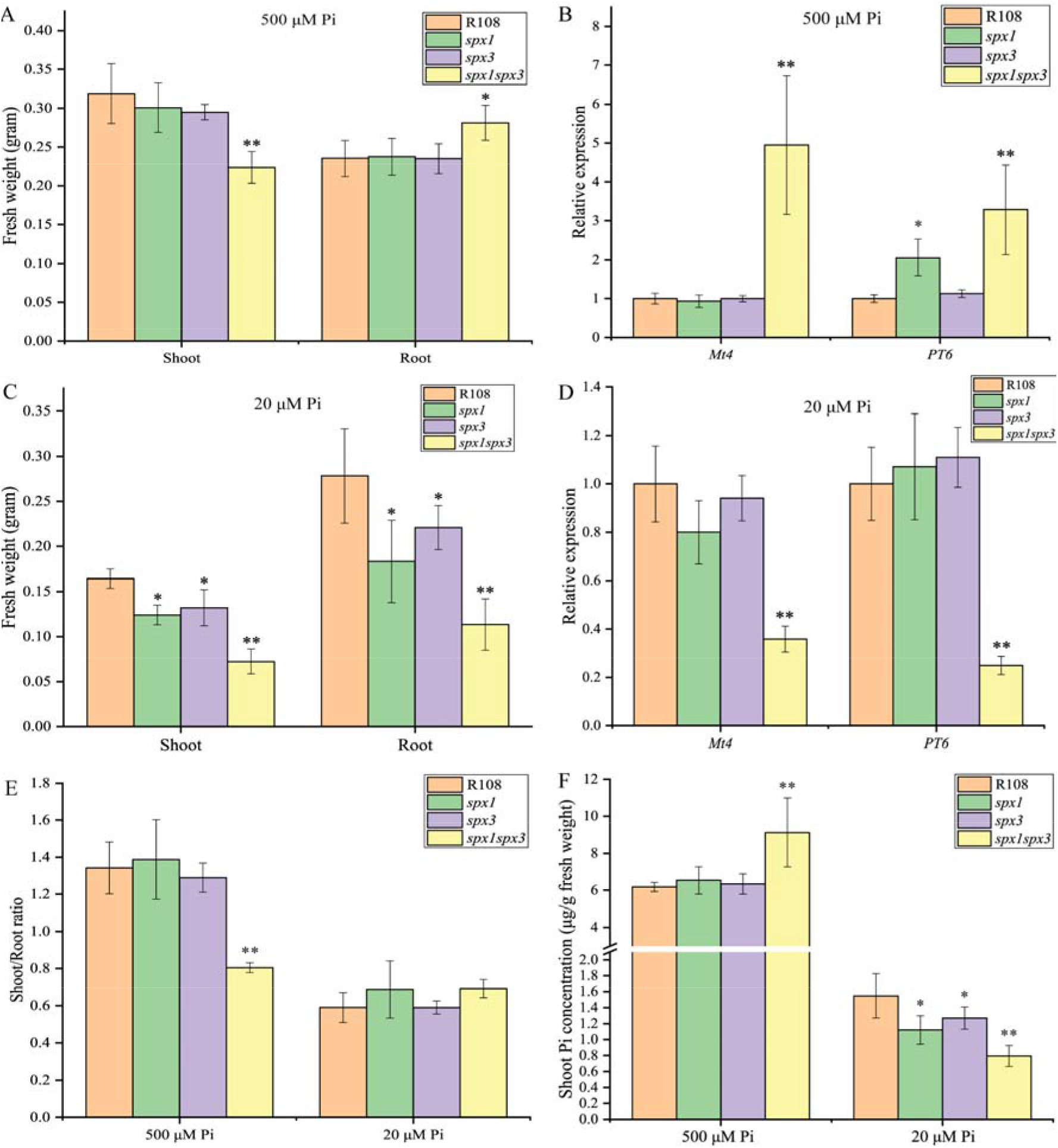
SPX1 and SPX3 regulate Pi homeostasis. (A) Fresh weight of wild type R108, *spx1*, *spx3* and *spx1spx3* plants grown for 3 weeks at 500 μM Pi. (B) Relative expression of *Mt4* and *PT6* in root samples shown in (A) as determined by qPCR. *MtEF1* is used as reference gene. (C) Fresh weight of wild type R108, *spx1*, *spx3* and *spx1spx3* plants grown for 3 weeks at 20 μM Pi condition. (D) Relative expression of *Mt4* and *PT6* in the root samples shown in (C). (E) Shoot-to-Root ratio of R108, *spx1*, *spx3* and *spx1spx3* plants shown in (A, C). (F) Shoot cellular Pi concentrations in R108, *spx1*, *spx3* and *spx1spx3* plants from (A, C). All values represent mean ± standard error of 5 replicate plants. Data significantly different from the corresponding R108 wild-type controls are indicated * P < 0.05; ** P < 0.01 (Student’s t-test).

At low (20 μM) Pi conditions the fresh weight of the *spx1* and *spx3* single mutants was significantly lower than wild-type R108 plants, and the *spx1spx3* double mutant showed an additive effect (Fig. 2C-F, S6B). The *spx1spx3* double mutant also showed a slight but significant higher shoot/root fresh weight ratio (Fig. 2E). The leaves of the double mutant further showed accumulation of anthocyanins in the leaves indicative of Pi starvation stress (Fig. S6B). Consistently, the PSI genes *Mt4* and *PT6* were lower expressed in the double mutant compared to the wild type (Fig. 2D), while overexpression of *SPX1/3* enhanced the expression of *Mt4* and *PT6* at low Pi conditions (Fig. 3E). Shoot Pi concentration in single mutants and double mutant were all significantly lower compared to the wild type (Fig. 2F). These results suggest that SPX1 and SPX3 also play a positive role in the PSR response under limiting (20 μM) Pi conditions.

**Fig. 3.**
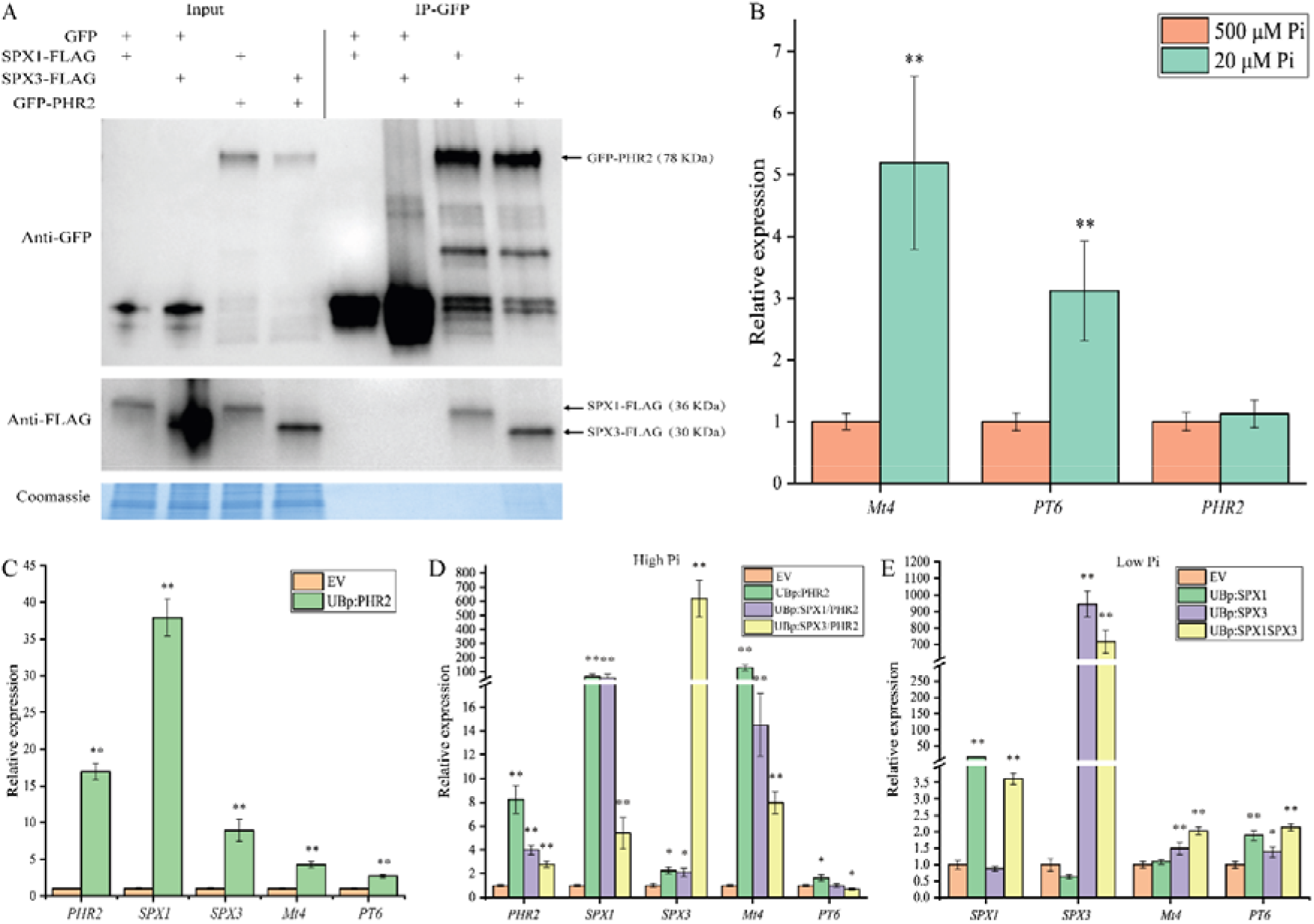
SPX1 and SPX3 interact with PHR2 to regulate phosphate homeostasis. (A) Western blot of co-immunoprecipitation samples showing SPX1 and SPX3 interaction with PHR2. FLAG-tagged SPX1 or SPX3 were co-expressed with free GFP or GFP-tagged PHR2 in *Nicotiana* leaves. Immunoprecipitation (IP) of GFP-tagged proteins shows the co-IP of the FLAG-tagged SPX proteins. Coomassie brilliant blue staining shows total protein levels as loading control. (B) qPCR analysis showing phosphate starvation induced *Mt4* and *PT6* expression, but not of *MtPHR2* expression in Medicago roots. Data represent mean ± standard error of 3 replicate plants. Data significantly different from 500 μM Pi conditions are indicated ** P < 0.01 (Student’s t-test). (C) qPCR analysis showing that overexpression of *MtPHR2* (*LjUBp::PHR2*) induced *SPX1*, *SPX3*, *Mt4* and *PT6* expression in transgenic roots grown for 4 days at low Pi conditions. (D) qRT-PCR results showing that overexpression (using the *LjUB1* promoter) of *SPX1* or *SPX3* together with *PHR2* in high Pi conditions induced *Mt4* and *PT6* expression less than overexpression of *PHR2* alone. (E) qPCR results showing that overexpression of *SPX1/3* at low Pi conditions induced *Mt4* and *PT6* expression. All values in (C-E) represent mean ± standard error of 3 independently transformed roots. Data significantly different from the corresponding EV transformed controls are indicated * P < 0.05; ** P < 0.01 (Student’s t-test). *MtEF1* was used as reference gene for normalization.

Overall, these results indicate that SPX1 and SPX3 are able to enhance the Pi starvation response at low Pi conditions and inhibit the Pi starvation response at high Pi conditions.

### SPX1 and SPX3 interact with PHR2

In Arabidopsis and rice, SPX proteins have been reported to interact with PHR and inhibit its activity under high Pi conditions (Wang et al., 2014; Puga et al., 2014). Therefore, we checked whether SPX1 and SPX3 could also interact with Medicago PHR homologs. Phylogenetic analyses indicated the presence of three PHR-like proteins in Medicago (Fig. S7). Co-immunoprecipitation analyses of GFP-tagged PHR with FLAG-tagged SPX1/3 proteins expressed in Nicotiana leaves revealed a clear interaction of both SPX1 and SPX3 with MtPHR2 (Medtr1g080330; hereafter PHR2 Fig. 3A). No significant interaction was found for the other two Medicago PHR-like proteins.

To study whether PHR2 is indeed involved in the phosphate starvation response, we overexpressed *PHR2* using the *LjUB1* promoter in Medicago roots and analyzed the effect on the expression of PSI genes *Mt4 PT6*(Mbodj et al., 2018; Hu et al., 2019). Both *Mt4* and *PT6* are strongly induced under Pi limiting conditions and also show induced expression upon overexpression of *PHR2* (Fig. 3B, C). *PHR2* itself was not regulated in a Pi-dependent manner at the transcriptional level (Fig. 3B), in analogy to its homologs *AtPHR1* (Bustos et al., 2010) and *OsPHR2* (Zhou et al., 2008).

To determine whether the observed Pi related phenotypes are due to a negative regulation of PHR2 activity by SPX1 and SPX3 we overexpressed of *SPX1&3* together with *PHR2* using the *LjUB1* promoter in Medicago roots. This showed that overexpression of *SPX1* or *SPX3* could indeed inhibit the induction of *Mt4* and *PT6* by PHR2 under high Pi conditions (Fig. 3D). We noticed that the (over)expression level of *PHR2* was ~ 2x lower for the co-expression constructs containing *SPX1* and *SPX3* compared to overexpression of PHR2 alone. This is likely a result of the expression construct rather than an effect of SPX1/3 on the activity of the *LjUB1* promoter. The much stronger (> 40x) reduced *Mt4* levels upon co-expression of *SPX1/3* support the inhibitory effect of SPX1/3 on the activity of PHR2 when sufficient Pi levels are reached.

### SPX1 and SPX3 regulate AM colonization and arbuscule degeneration

Next, we examined the role of SPX1 and SPX3 in the interaction with AM fungi. Three weeks after inoculation with *R. irregularis* spores, mycorrhization was quantified in the *spx1* and *spx3* single mutants, the *spx1spx3* double mutant and R108 wild-type controls using the magnified intersect method (McGONIGLE et al., 1990). Eight plants were used as replicates for each line. Compared to R108, single mutant and double mutant plants all showed significantly lower root colonization levels and arbuscule abundance (Fig. 4A). Transcript levels of *RiEF* and *PT4*, molecular markers for respectively fungal colonization and arbuscule abundance, confirmed the lower colonization levels in the mutants (Fig. 4B).

**Fig. 4.**
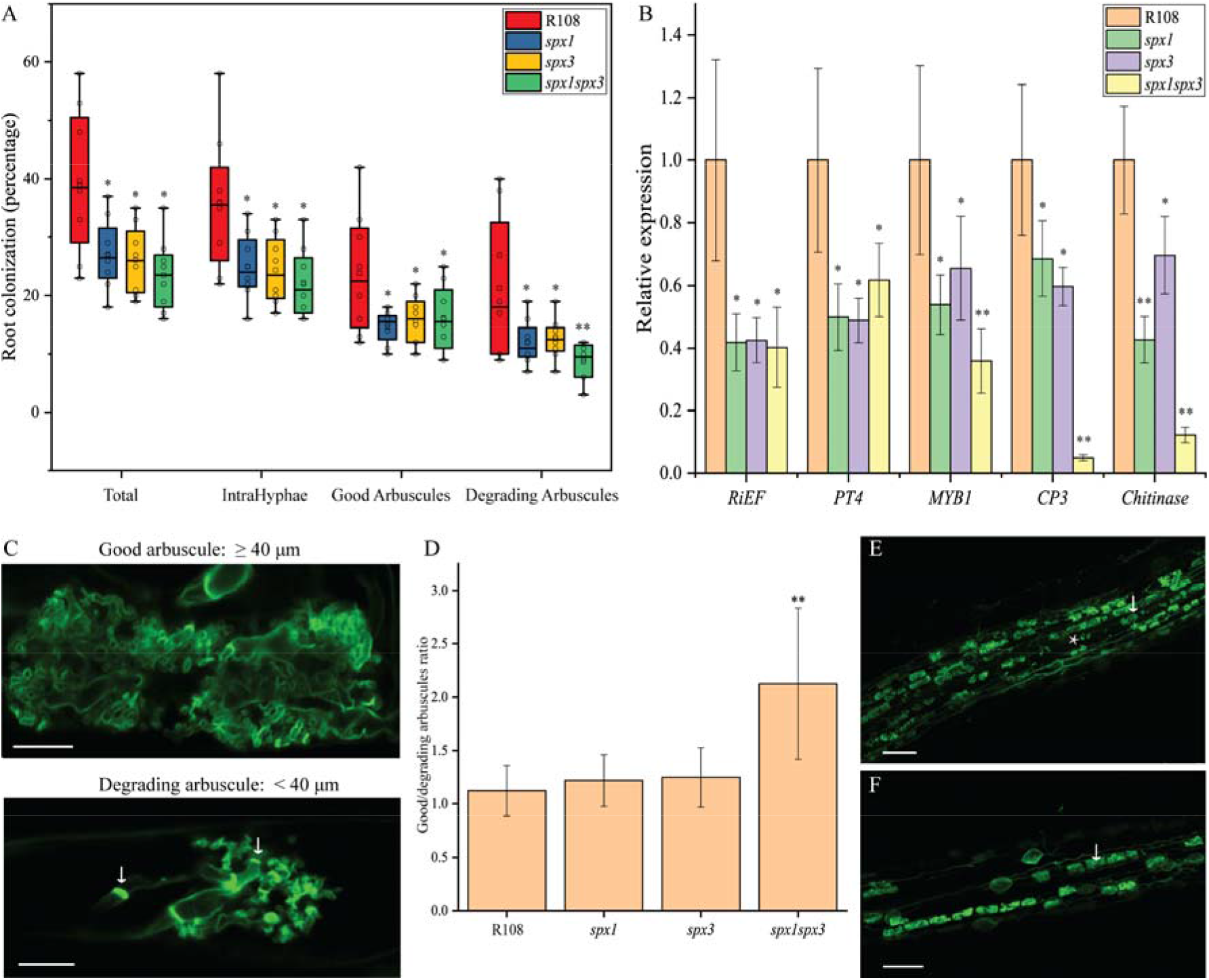
SPX1 and SPX3 regulate AM colonization and arbuscule degeneration. (A) Quantification of mycorrhization levels in wild type R108, *spx1*, *spx3* and *spx1spx3* 3 weeks post inoculation with *R. irregularis*. 8 independently transformed roots were used as replicas for each sample. Quantification was performed using the magnified intersections method (McGONIGLE et al., 1990). Good arbuscule: larger or equal to 40 μm; Degrading arbuscules: smaller than 40 μm with septa (white arrow in C). Data significantly different from R108 wild-type controls are indicated * P < 0.05; ** P < 0.01 (Student’s t-test). (B) Expression levels of *RiEF*, *PT4*, *CP3*, *Chitinase* and *MYB1* in root samples from (A) determined by qPCR analyses. *MtEF1* was used as internal reference. Values represent mean ± standard error. Data significantly different from R108 wild-type controls are indicated * P < 0.05; ** P < 0.01 (Student’s t-test). (C) Representative images of good arbuscule and degrading arbuscules in WGA-Alexa488 stained roots 3 weeks post-inoculation. Arrow points to septa. Scale bar = 10 μm. (D) Good-to-degrading arbuscule ratio of R108, *spx1*, *spx3* and *spx1spx3* mycorrhizal samples from (A). Values represent mean ± standard error of 8 independently transformed roots. Data significantly different from R108 wild-type controls are indicated ** P < 0.01 (Student’s t-test). (E) and (F) Representative images of WGA-Alexa488 stained *R. irregularis* in R108 and *spx1spx3*. White arrow marks a good arbuscule. Asterisk marks a degrading arbuscule. Scale bar = 100 μm.

Because *SPX1* and *SPX3* are specifically expressed in arbuscule-containing cells in mycorrhized roots, arbuscule morphology was quantified in more detail. We defined arbuscules larger or equal to 40 μm as “good” arbuscules, and arbuscules smaller than 40 μm with typical features of degradation, including visible septa, as “degrading” arbuscules (Fig. 4C). Interestingly, there were significantly less degrading arbuscules in the *spx1spx3* double mutant compared to R108 wild-type plants, resulting in a much higher good/degrading arbuscule ratio (Fig. 4A, E, F). *spx1* and *spx3* single mutant showed a similar good/degrading arbuscule ratio as wild-type R108 (Fig. 4A), suggesting a redundant role for SPX1 and SPX3 in the regulation of arbuscule degradation.

Arbuscule degradation in Medicago is regulated by the MYB1 transcription factor, which controls the expression of hydrolase genes such as *Cysteine Protease 3* (*CP3*) and *Chitinase* (Floss et al., 2017). qRT-PCR analyses showed a strongly impaired expression of these hydrolase genes in the *spx1spx3* double mutant (Fig. 4B). Similar phenotypes were observed upon knock-down of both *SPX1* and *SPX3* by RNA interference (Fig. S8A, B). To further confirm that the phenotype was indeed caused by a mutation in *SPX1* and *SPX3*, we complemented the *spx1spx3* double mutant by driving *SPX1* and *SPX3* expression from by their native promoters in *A. rhizogenes* transformed roots. This indeed complemented the mycorrhization levels, arbuscule abundance and marker gene expression to wild-type levels (Fig. S8C, D), showing that the phenotypes are not caused by background insertions/mutation in the mutant lines.

Overall, these results indicate that both SPX1 and SPX3 positively regulate AM colonization levels and redundantly regulate arbuscule degradation.

### SPX1 and SPX3 likely control AM colonization by regulating strigolactone levels

To explain the positive role of SPX1 and SPX3 in mycorrhizal colonization, we found that the expression of a key gene required for strigolactone biosynthesis, *MtDWARF27* (*D27*; (Hao et al., 2009; Liu et al., 2011) was not (or much lower) induced upon Pi starvation in *spx1* and *spx3* single mutants compared to wild-type plants (Fig. 5A). No additive effect on *D27* expression was observed in the *spx1spx3* double mutant. Under low phosphate conditions, strigolactone levels increase drastically in several species and this induction in strigolactone biosynthesis correlates with an increased *D27* expression under this condition (Liu et al., 2011) (Fig. 5A). *D27* expression was induced when *SPX1* and *SPX3* were overexpressed together at low Pi conditions using the *LjUB1* promoter in *A. rhizogenes* transformed roots (Fig. S9A). At high Pi conditions *D27* expression did not appear to be affected (Fig. S9B). This revealed a positive effect of SPX1 and SPX3 on *D27* expression at low Pi conditions, and thereby possibly on strigolactone levels. Reduced strigolactone levels could explain the lower colonization levels observed in the mutants as these are key signal molecules that induce growth and branching in AM fungi (Besserer et al., 2006; Tsuzuki et al., 2016). Unfortunately, we were unable to detect known strigolactone levels in the R108 genetic background. This may be caused by sub-detection levels or as yet unknown strigolactone derivates in R108. As an alternative, we used an AM hyphal branching assay as proxy for strigolactone levels in root exudates (Besserer et al., 2006, 2008). This demonstrated that exudates collected from the *spx1/3* mutant roots, grown under Pi limiting conditions, were much less able to induce *R. irregularis* branching compared to wild-type R108 root exudates (Fig. 5B, S8C). Together these data strongly suggests that strigolactone levels are reduced in the *spx* mutants, although an additional effect on other root exudates cannot be ruled out.

**Fig. 5.**
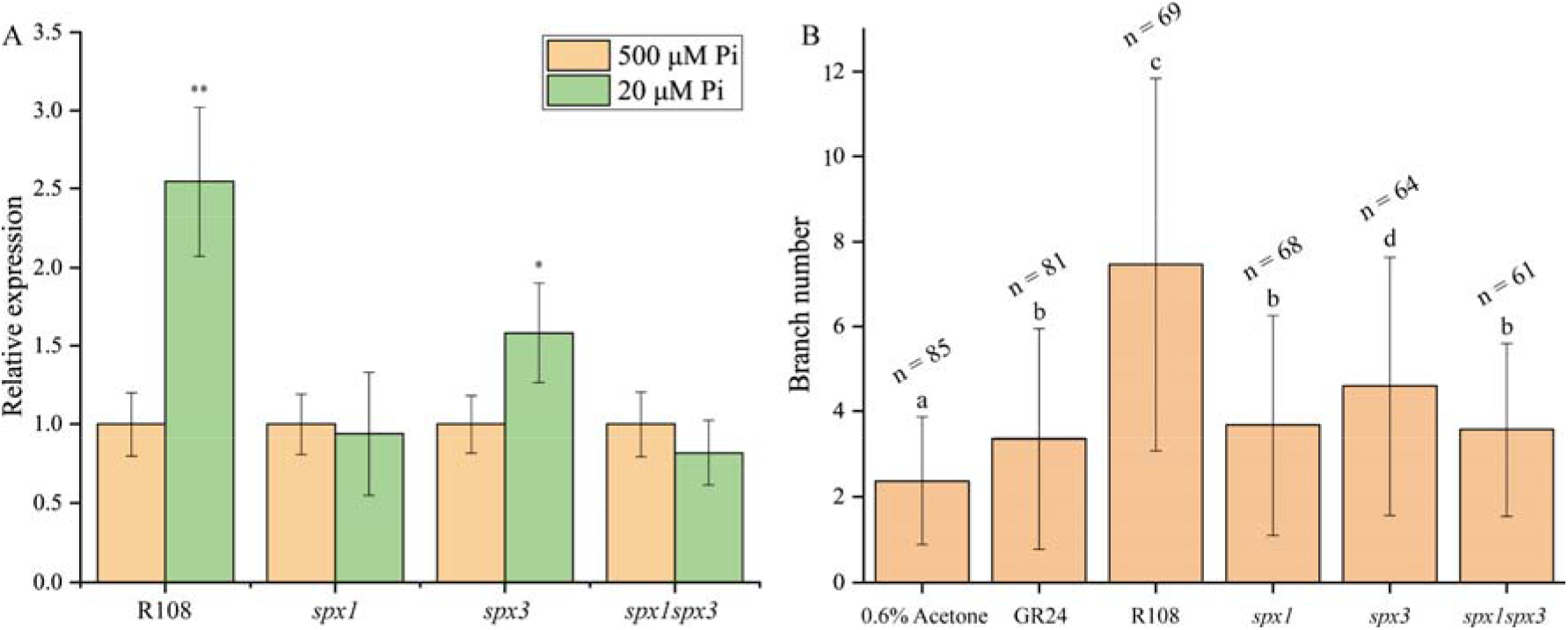
SPX1 and SPX3 regulate expression of strigolactone biosynthesis gene *D27*. (A) The induction of *D27* expression in low Pi conditions is impaired in *spx1*, *spx3*, and *spx1spx3* mutants. Values represent mean ± standard error of 3 individual plant replicates. Data significantly different from corresponding controls are indicated * P < 0.05; ** P < 0.01 (Student’s t-test). (B) *R. irregularis* spores treated with root exudates of R108, *spx1*, *spx3* and *spx1spx3* for 8 days. 0.01 μM GR24 and 0.6% acetone were respectively used as positive and negative controls. Hyphal branch numbers in spores treated with root exudates of *spx1*, *spx3* and *spx1spx3* mutants are significantly lower compared to spores treated with R108 root exudates. n indicates the number of spores counted. Error bars indicate standard error, different letters indicate significant difference between treatments (one-way ANOVA, P < 0.05).

### Overexpression of SPX1 and SPX3 increases AM colonization and arbuscule degradation

The mutant analyses showed that SPX1 and SPX3 redundantly regulate arbuscule degradation. To further study this we overexpressed *SPX1, SPX3,* and *MtSPX1;MtSPX3* together under the control of the *LjUB1* promoter. This resulted in significantly increased colonization levels in individual *LjUBp:SPX1* and *LjUBp:SPX3* transgenic roots as well as double transgenic lines compared to empty vector control roots 3 weeks post inoculation (Fig. 6A). Furthermore, the ratio of good/degrading arbuscule was significantly lower in the *SPX1/3* overexpression roots indicating a premature degradation of arbuscules (Fig. 6A-C). qRT-PCR analyses confirmed the observed phenotypes (Fig. S10A). *RiEF* was higher expressed in *SPX1* and *SPX3* overexpression roots compared to empty vector (EV) controls expressed roots. The expression level of *PT4* as marker for healthy arbuscules was similar between EV control roots and the *SPX* overexpressing roots. However, arbuscule degradation related genes *CP3* and *Chitinase* were significantly enhanced in the *SPX* overexpressing roots. We observed a differential effect of *SPX1* and *SPX3* overexpression on *MYB1* expression, with only *SPX3* overexpression causing increased *MYB1* expression (Fig. S10A).

**Fig. 6.**
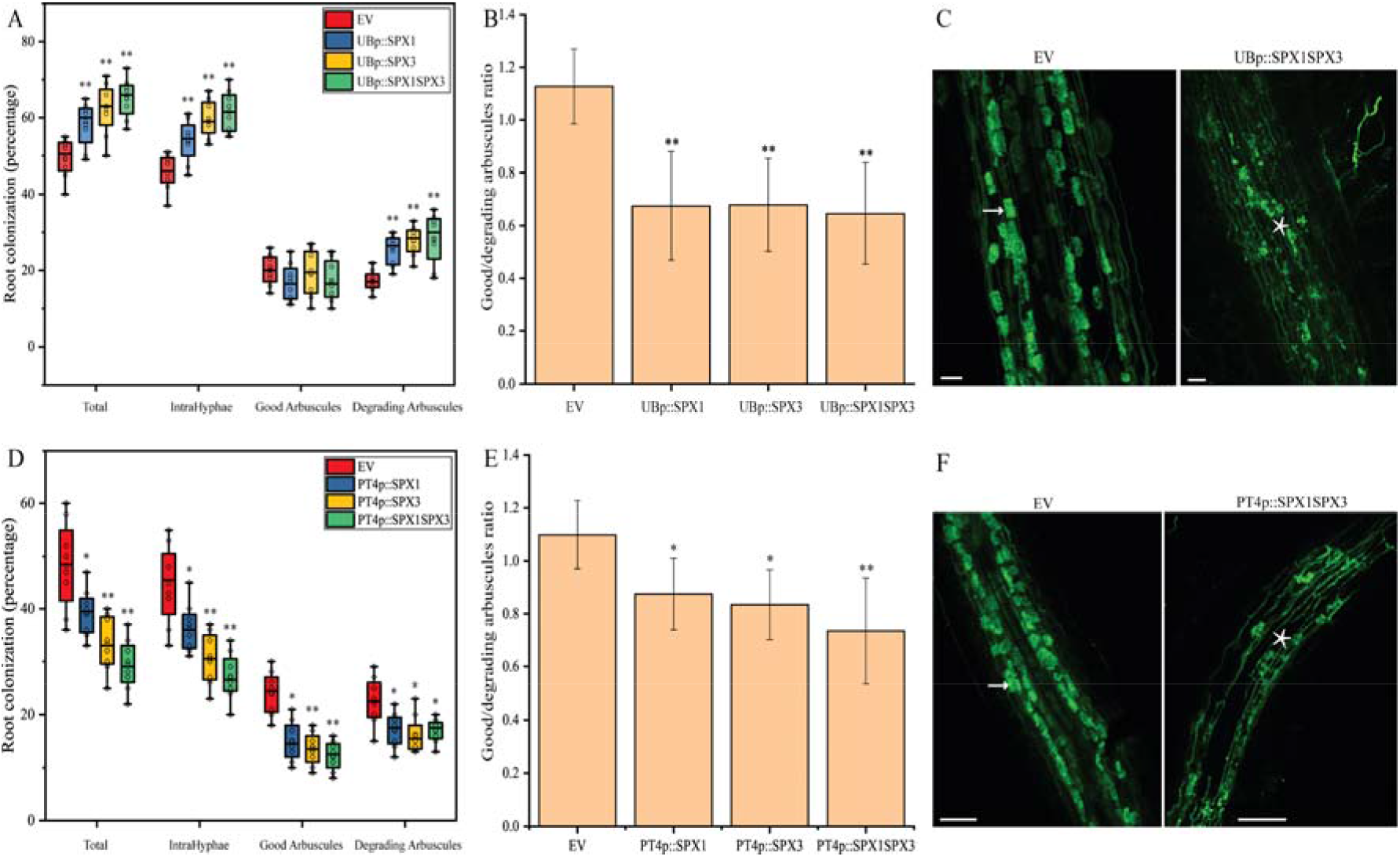
Overexpression of *SPX1/3* under the control of the *LjUbiquitin* or *MtPT4* promoter increased AM colonization and/or arbuscule degradation. (A) Quantification of mycorrhization level in *M. truncatula* roots expressing EV, *UBp::SPX1*, *UBp::SPX3*, and *UBp::SPX1-UBp::SPX3* 3 weeks post inoculation with *R. irregularis*. 8 independently transformed plants were used as replicas for each sample. Quantification was performed using the magnified intersections method (McGONIGLE et al., 1990). (B) Good-to-Degrading arbuscule ratio of EV, *UBp::SPX1*, *UBp::SPX3* and *UBp::SPX1-UBp::SPX3* mycorrhizal samples from (A). Values represent mean ± standard deviation of 8 independently transformed roots. (C) Representative confocal images of WGA-Alexa488 stained *R. irregularis* in EV and *UBp::SPX1-UBp::SPX3* roots. White arrow marks a good arbuscule. Asterisk marks a degrading arbuscule. Scale bar = 80 μm. (D) Quantification of mycorrhization level in *M. truncatula* roots expressing EV, *PT4p::SPX1*, *PT4p::SPX3* and *PT4p::SPX1-PT4p::SPX3* 3 weeks post inoculation with *R. irregularis*. 8 independently transformed plants were used as replicas for each sample. Quantification was performed using the magnified intersections method (McGONIGLE et al., 1990). (E) Good-to-Degrading arbuscule ratio of EV, *PT4p::SPX1*, *PT4p::SPX3* and *PT4p::SPX1-PT4p::SPX3* mycorrhizal samples from (D). Values represent mean ± standard deviation of 8 independently transformed roots. (F) Representative confocal images of WGA-Alexa488 stained *R. irregularis* in EV and *PT4p::SPX1-PT4p::SPX3* roots. White arrow marks a good arbuscule. Asterisk marks a degrading arbuscule. Scale bar = 100 μm. All data in (A, B, D, E) significantly different from the corresponding EV controls are indicated * P < 0.05; ** P < 0.01 (Student’s t-test).

To separate the arbuscule-specific effects of SPX1/3 from non-symbiotic roles, we overexpressed *SPX1*, *SPX3,* or both, using the arbuscule-specific *PT4* promoter. Interestingly, compared to EV transformed roots decreased colonization levels were observed in all *SPX* overexpressing roots when expressed from the *PT4* promoter (Fig. 6D). This coincided with decreased arbuscule abundance and an increased ratio of degrading arbuscule compared to good arbuscule classes (Fig. 6D-F). Transcript levels of the markers for fungal biomass (*RiEF*), healthy arbuscule abundance (*PT4*) and arbuscule degradation (*MYB1, CP3* and *Chitinase*) all confirmed the visual phenotyping results (Fig. S10B). Because the colonization levels in *SPX* overexpressing roots were much lower than EV control roots and because *SPX1* and *SPX3* are normally highly induced in arbuscule-containing cells the overall *SPX1/3* expression levels did not exceed those in the EV controls (Fig. 1B, 6D, S10B). Overall, these results further strengthen the role for SPX1/3 in regulating AM colonization and arbuscule degradation.

### SPX1 and SPX3 in relation to known transcriptional regulators of arbuscule degradation

The MYB1 transcription factor was reported to be required for arbuscule degradation when the fungus does not provide sufficient nutrients. Knock-down of *MYB1* could rescue the premature arbuscule degradation phenotype observed in *pt4* mutants that lack a functional symbiotic PT4 phosphate transporter (Floss et al., 2017). To study whether SPX1/3 can also rescue the *pt4* phenotype, we knocked down both *SPX1* and *SPX3* using RNAi in the *pt4-1* mutant background (Javot et al., 2007). This resulted in strongly reduced mycorrhization levels in *EFp:SPX1-SPX3-rnai* transformed *pt4-1* mutant roots compared to EV controls 3 week after inoculation (Fig. S11A). However, the ratio of “Good” to “degrading” arbuscules was not significantly different between SPX1/3 RNAi-*pt4* and EV-*pt4* samples (Fig. S11A, B). qPCR analyses further confirmed the observed phenotypes (Fig. S11C). This suggests that *SPX1/3* function is not essential for arbuscule degradation in this setting.

It was further shown that overexpression of *MYB1* can trigger the premature degradation of arbuscules (Floss et al., 2017). To position the action of SPX1/3 on arbuscule development in relation to MYB1, we overexpressed *MYB1* under the control of the constitutive CaMV35S promoter in the *spx1spx3* double mutant and checked whether *CP3* and *Chitinase* could still be activated. Compared to EV transformed roots, *CP3* and *Chitinase* expression were significantly induced by *MYB1* in the *spx1spx3* double mutant 3 weeks post inoculation and lower colonization levels, with less *RiEF* and *PT4* expression, and signs of premature arbuscule degradation were detected (Fig. S11D, S12). This indicates that SPX1/3 functions either upstream or parallel of MYB1 to control arbuscule turnover. To distinguish between these possibilities, we overexpressed both *SPX1* and *SPX3* together and studied the effect on *MYB1* expression and its target genes under non-symbiotic conditions. At both high and low Pi conditions, overexpression of *SPX1* and *SPX3* (*LjUB1p:SPX1-LjUB1p:SPX3*) did not (significantly) affect *MYB1* expression (Fig. S11E, F), which suggests that SPX1 and SPX3 likely do not function upstream of MYB1 to directly control its expression. The expression of the hydrolase-encoding gene *CP3* was also not induced upon overexpression of *SPX1&3* and the expression of *Chitinase* was only weakly affected at low Pi conditions (Fig. S11E, F). The lack of/weak effect on *CP3* and *Chitinase* expression may be caused by the low expression levels of MYB1 under non-symbiotic conditions.

To investigate a link between SPX1/3 and the MYB1 transcription complex further we studied whether SPX1 and SPX3 interact with MYB1 or its interacting partners, the GRAS transcription factors NSP1 and DELLA, which are required for MYB1’s ability to induce *CP3* and *Chitinase* (Floss et al., 2017). Co-immunoprecipitation analyses of FLAG-tagged SPX1 and SPX3 expressed in *Nicotiana benthamiana* leaves together with either GFP-MYB1, NSP1-GFP or GFP-DELLA1-Δ18 (a dominant version of DELLA1 (Floss et al., 2013) failed to detect a significant interaction between any of these proteins. In contrast, we did detect the interaction between MYB1 and NSP1 as reported by Floss et al. (2017). Also yeast-two hybrid analyses did not detect a significant interaction between SPX1/3-AD and MYB-BD or DELLA1-Δ18-BD (data not shown). Expression of NSP1-BD caused the autoactivation of reporters in our yeast-two-hybrid system preventing the observation of a potential interaction.

Our inability to detect significant interactions with the currently known regulators of arbuscule degradation, suggest that other regulators remain to be identified. Based on the results obtained in this study we propose the following model (Fig. 7), where SPX1 and SPX3 act as negative regulators of a yet to identify negative regulator of MYB1. This negative regulation of MYB1 activity likely requires sufficient Pi conditions in the arbuscule containing cells. When cells have acquired sufficient Pi SPX1/3 will ensure the timely degradation of the arbuscules.

**Fig. 7.**
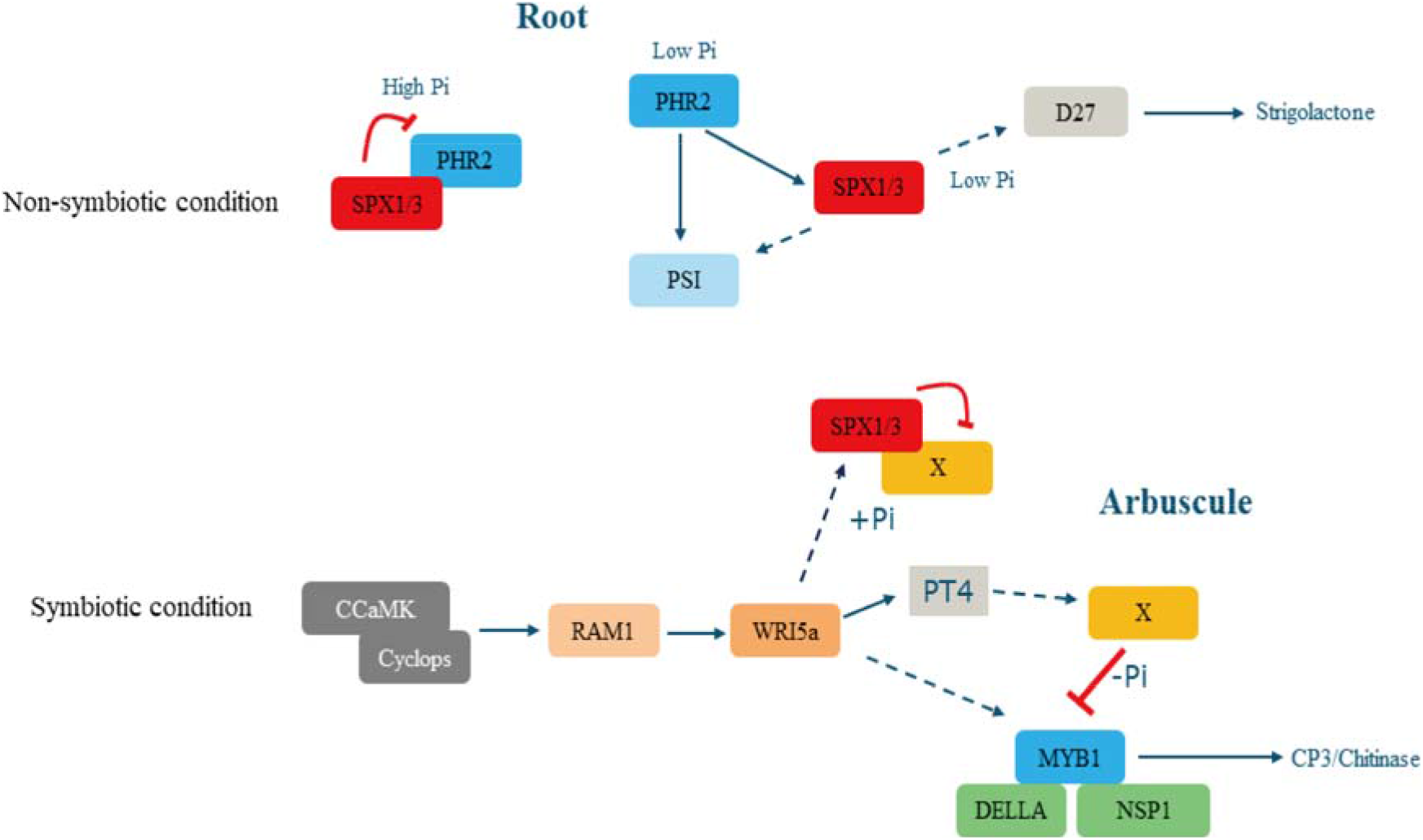
Proposed model for SPX1/3 function. In high Pi conditions, SPX1 and SPX3 interact with PHR2, inhibiting PHR2-induced PSI gene expression. In low Pi conditions, PHR2 induces phosphate starvation induced (PSI) genes as well as *SPX1/3* expression. SPX1 and SPX3 play a positive role in the expression of PSI genes (*Mt4* and *PT6*) and *D27*. *D27* is a key gene involved in strigolactone biosynthesis. Strigolactones act as signaling molecules to enhance the growth and metabolism of AM fungi. In symbiotic conditions, SPX1 and SPX3 redundantly regulate arbuscule degeneration likely through an yet-to-be-identified factor X, which negatively regulate MYB1 activity. *SPX1* and *SPX3* are induced by the RAM1-WRI5 transcriptional cascade that also induces *PT4* and *MYB1* expression. The activity of X is suppressed by binding of SPX1/3 in Pi-dependent manner. This relieves the inhibition of MYB1 activity that, together with DELLA and NSP1 transcription factors, induces arbuscule degradation genes such as *CP3* and *Chitinase* leading to arbuscule degradation. Red lines indicate Pi dependent negative regulation at the protein level. Solid arrows indicate known transcriptional induction. Dashed arrows indicate direct/indirect transcriptional regulation based on the data presented.

## Discussion

SPX proteins have emerged as key sensors and signaling regulators of cellular phosphate status in plants (Wild et al., 2016; Wang et al., 2014; Puga et al., 2014; Shi et al., 2014; Hu et al., 2019). Here we show that the Medicago single SPX-domain proteins SPX1 and SPX3 not only regulate Pi homeostasis under non-symbiotic conditions, but also regulate root colonization and arbuscule degradation during AM symbiosis. This offers important novel insight into the Pi-dependent regulation of this agriculturally and ecologically important symbiosis.

Under non-symbiotic conditions, SPX1 and SPX3 control Pi homeostasis in part through the regulation of PHR2 activity. In analogy to the situation in *Arabidopsis* and rice, phosphate starvation leads to the activation of PHR activity to control transcriptional responses. Among the targets of PHR2 are the *SPX1/3* genes. Both SPX1 and SPX3 can bind PHR2 at high Pi conditions and negatively affect the PSR response to prevent overaccumulation of Pi. We show that SPX1 and SPX3 also control the induction of strigolactone biosynthesis gene *D27* (Liu et al., 2011) under Pi-limiting conditions. This suggests that SPX1 and SPX3 play an additional positive role in the transcription of Pi-starvation induced genes under low Pi conditions. This is further supported by the observation that SPX1 and SPX3 play a positive role in the PSR response at low Pi conditions, in addition to their negative regulation of PSR at high Pi conditions. Such additional, possibly PHR-independent roles, have also been suggested for OsSPX3 and OsSPX5 (Shi et al., 2014). Since D27 plays an essential role in the biosynthesis of strigolactones (Hao et al., 2009), which activate AM fungi, the role of SPX1 and SPX3 in its induction could explain the lower colonization levels observed in the *spx1* and *spx3* mutants. This hypothesis is strongly supported by the reduced stimulatory effect of root exudates from the *spx1* and *spx3* mutant on AM hyphal branching compared to the R108 wild type. However, the current inability to measure strigolactone levels in the R108 genetic background prevented us from confirming this directly. It would also explain the stimulatory effect of overexpression of SPX1/3 using the constitutive *LjUB1* promoter on fungal colonization levels (Fig. 6a). In contrast, overexpression of *SPX1/3* under the control of the arbuscule-specific *PT4* promoter leads to a decrease in fungal colonization (Fig. 6d). The arbuscule-specific expression likely prevents the stimulatory effect of SPX1/3 on strigolactone (or additional metabolites) exudation while still reducing the level of functional arbuscules leading to overall lower colonization levels.

The induction of *SPX1* and *SPX3* upon Pi starvation or *PHR2* overexpression fits with the presence of the P1BS *cis*-regulatory element in the promoters of these genes. However, after establishment of the AM symbiosis, the expression of both *SPX1* and *SPX3* becomes restricted from an ubiquitous expression pattern to specific expression in the arbuscule-containing cells. This suggests that activity of PHR in the non-colonized root cortical and epidermal cells is inhibited upon a functional AM symbiosis in a non-cell autonomous manner. It has been proposed that AM fungi may interfere with the direct phosphate uptake of plants (Smith et al., 2004; Christophersen et al., 2009; Yang et al., 2012; Wang et al., 2020), although the mechanisms for this are still unknown. Another possibility is a more systemic regulation of the phosphate starvation response through hormonal or peptide signaling as the plant is obtaining Pi from the fungus (Müller and Harrison, 2019). Although we cannot pinpoint at what time after initiation of the symbiosis, or level of colonization, the shift in expression exactly occurs based on our analyses, the fact that we hardly see expression in non-colonized root cells, nor in non-transgenic roots on the same composite plants, already after 3 weeks of inoculation suggests a rather fast systemic regulation.

It is currently not known whether PHR2 is active in arbuscule-containing cells, as contrasting roles for the P1BS element in the expression of symbiotic phosphate transporters have been reported (Chen et al., 2011; Lota et al., 2013). However, the presence of SPX1/3 would be expected to suppress PHR2 activity as it does under non-symbiotic conditions. Instead, the induction of *SPX1* and *SPX3* in arbuscule-containing cells correlates with the presence of multiple AW-boxes and CTTC elements in the promoters of both genes. These *cis*-regulatory elements are found in many genes that are induced in arbuscule-containing cells, and are bound by WRINKLED1-like TFs that are in turn activated by the key GRAS transcription factor RAM1 that controls arbuscule formation (Jiang et al., 2018; Xue et al., 2018; Limpens and Geurts, 2018; Pimprikar et al., 2018). A link between RAM1, WRI5a and *SPX3* expression is further supported by the lack of *SPX3* induction in the *ram1* mutant (table S3) (Luginbuehl et al., 2017). However, this hypothesis still requires further experimental confirmation.

Since Pi levels are most likely not limiting in cells containing active arbuscules, the impaired induction of *CP3* and *Chitinase* and associated arbuscule degradation in the *spx1spx3* double mutant argues for the involvement of SPX1/3 interacting proteins other than PHR2. The observation that *MYB1* overexpression was able to trigger premature arbuscule degradation in the *spx1spx3* double mutant indicates that MYB1 can bypass SPX1/3 to induce arbuscule degradation. We were unable to detect a significant interaction between SPX1 or SPX3 with the currently known regulators of arbuscule degradation, MYB1 and its interacting partners NSP1 and DELLA (Floss et al., 2017). This leaves us with the question how SPX1 and SPX3 might control arbuscule degradation? Based on our findings we propose that SPX1 and SPX3 negatively regulate a yet unknown negative regulator of MYB1 activity in a Pi-dependent manner (Fig. 7). *MYB1* expression is already strongly induced at an early stage of arbuscule formation, most likely due to the RAM1-WRI5 transcriptional cascade. However this induction does not lead to the degradation of the arbuscules (Floss et al., 2017). Therefore, it seems that the activity of MYB1 is suppressed as long as the fungus provides (sufficient) Pi to the host cell. Ectopic overexpression of *MYB1* can trigger the premature degradation of the arbuscules suggesting that the proposed negative regulator is not expressed yet or that high MYB1 levels can override the negative regulation. SPX1 and SPX3 are predicted to sense the amount of Pi proved by the fungus in the arbuscule-containing cells. If Pi levels exceed the demand SPX1 and SPX3 can redundantly inhibit the negative regulation of MYB1 activity thereby terminating the arbuscule.

It is tempting to speculate that SPX proteins may also be involved in the cross-talk with other nutrients supplied by the fungus, such as nitrogen. This suggestion is based on the involvement of OsSPX4 in nitrate signaling to activate the phosphate starvation response (Hu et al., 2019) and the reported nitrogen-control of arbuscule development (Breuillin-Sessoms et al., 2015). For example, the premature arbuscule degradation in the *pt4* mutant was also suppressed when the plants were grown under N-limiting conditions (Breuillin-Sessoms et al., 2015). This was shown to depend on the ammonium transceptor AMT2;3. It will therefore be interesting to test whether SPX1 and SPX3 contribute to this nutrient crosstalk in Medicago.

In conclusion, we reveal novel roles for phosphate sensing SPX proteins in the regulation of AM symbiosis to enhance the phosphate acquisition efficiency of plants. SPX proteins show both a role in the initiation of the symbiosis, likely in part through their effect on the expression of the strigolactone biosynthesis gene *D27*, as well as a role in the termination of the symbiosis by controlling the degradation of arbuscules. The latter role can be essential to maintain the beneficial nature of the interaction. In nature plants are most often colonized by multiple different AM strains that can differ in the amount of nutrients they supply (Kiers et al., 2011). Therefore SPX proteins can provide a means to locally monitor whether a fungal partner provides sufficient nutritional benefits. Further unraveling how nutrient sensing and homeostasis are regulated during AM symbiosis will be pivotal to understand the ecological workings of this key symbiosis and how to best exploit it for more sustainable agriculture practices.

## Materials and methods

### Plant and fungal material

*Medicago truncatula* A17 and R108 seedlings were grown and transformed as described (Limpens et al., 2004). The *spx1* (NF13203) and *spx3* (NF4752) Tnt1-insertion lines were obtained from the Noble Research Institute (https://medicago-mutant.noble.org/mutant/index.php). Homozygous *spx1* and *spx3* mutants were identified by PCR and crossed to obtain the *spx1/spx3* double mutant. Primers used are listed in Table S1. Plants were grown in SC10 RayLeach cone-tainers (Stuewe and Sons, Canada) with premixed sand:clay (1:1 V/V) mixture and watered with 10 ml ½ Hoagland medium with 20 μM (low Pi) or 500 μM H_2_PO_4_ (high Pi) twice a week in a 16 h daylight chamber at 21℃, as described previously (Zeng et al., 2018). *Rhizophagus irregularis* DAOM197198 spores (Agronutrion, France) were washed through three layers of filter mesh (220 μm, 120 μm, 38 μm) before inoculation. 200 pores were placed ~2 cm below seedling roots.

### Phylogeny

SPX phylogenetic analysis was performed using the Geneious R11.0 software package (https://www.geneious.com). *Medicago*, *Arabidopsis* and rice SPX protein sequences were collected from PLAZA (https://bioinformatics.psb.ugent.be/plaza/) and aligned using MAFFT in Geneious R11.0. An unrooted SPX phylogenetic tree was generated using the neighbour-joining tree builder with 500 bootstraps.

### Constructs

Most constructs were made using the Golden gate cloning system (Engler et al., 2014) Constructs for RNAi were made via Gateway cloning and constructs for Y2H experiments were made using in fusion cloning (Takara, Japan). Primers used are listed in Table S1. Vectors used for cloning are listed in table S2. All newly made vectors were confirmed by Sanger sequencing.

### GUS histochemical analyses

To create the *MtSPX1p-GUS* and *MtSPX3p-GUS* constructs, a 1824 bp upstream of the *MtSPX1* (Medtr3g107393) ATG start codon and a 1260 bp upstream of the *MtSPX3* (Medtr0262s0060) ATG start codon was amplified by PCR from *M. truncatula* A17 genomic DNA as promoter, respectively. *MtSPX1p-GUS* and *MtSPX3p-GUS* constructs were introduced into Medicago plants using *Agrobacterium rhizogenes*-mediated root transformation (Limpens et al., 2004). GUS staining was done as described (An et al., 2019). Briefly, transgenic roots were harvested based on DsRed fluorescence (red fluorescent marker present in the constructs) and washed twice with PBS buffer for 10 min. Next, the roots were incubated in GUS reaction buffer (3% sucrose, 10mM EDTA, 2 mM potassium-ferrocyanide, 2 mM potassium-ferricyanide and 1 mg/ml X-Gluc in 100mM PBS, pH 7.0) for 30 min in vacuum, and then incubated at 37°C for 1 h. The stained roots were fixed in fixation buffer (5% glutaraldehyde in 100mM phosphate buffer, pH 7.2) for 2 h in vacuum at room temperature, followed by dehydrating with an ethanol series (20%, 30%, 50%, 70%, 90%, 100%) for 10 min each. Root segments were embedded in Technovit 7100 (Hereus-Kulzer, Germany) and cut into 8 μm longitudinal sections using a microtome (Leica RM2255) and stained with 0.1% Ruthenium Red for 5 min. Images were taken using an Leica DM5500 B microscope.

### Co-immunoprecipitation and western blotting

FLAG-tagged SPX1 and SPX3 and GFP-tagged PHR2 (Medtr1g080330) constructs were transiently expressed in *Nicotiana* leaves as described (Zeng et al., 2018). Total proteins were isolated using Co-IP buffer (10% glycerol, 50 mM Tris-Hcl pH=8.0, 150 mM NaCl, 1% Igepal CA 630, 1 mM PMSF, 20 μM MG132, 1 tablet protease inhibitor cocktail). GFP-Trap agarose beads (Chromotek) were used to immunoprecipitate GFP protein complexes. Western blot was performed as described (Bungard et al., 2010). 1:5000 diluted anti-GFP-HRP and anti-FLAG-HRP antibodies (Miltenyi biotec, USA) were used for detection.

### RNA interference

A *SPX1SPX3* hairpin construct was generated targeting both *SPX1* and *SPX3* mRNA using the Gateway system (Invitrogen, USA). 534 bp of *SPX1* and 489 bp of *SPX3* mRNA sequence were combined together by overlap PCR and the resulting 1023 bp sequence was cloned into the pENTR/D-TOPO entry vector. Primers used are listed in table S1. the. Subsequently, the modified pK7GWIWG2(II)-AtEF1 RR vector (Zeng et al., 2020) was used for an LR reaction to get the final hairpin silencing construct.

### Pi concentration measurement

Shoots inorganic Pi concentration was measured as described (Zhou et al., 2008). Briefly, Medicago shoots were crushed to a fine powder in liquid nitrogen, and rigorously vortexed for 1 min in 1 mL 10% (w/v) of perchloric acid. The homogenate was diluted 10 times by 5% (w/v) perchloric acid and then put on ice for 30 min, followed by centrifugation at 10,000g for 10 min at 4°C to collect the supernatant. The molybdenum blue method was used to measure Pi content in the supernatant. To prepare the molybdenum blue solution, 6 mL solution A (0.4% ammonium molybdate dissolved in 0.5 M H_2_SO_4_) was mixed with 1 mL 10% ascorbic acid. 2 mL of molybdenum blue solution was added to 1 mL of the sample supernatant, and incubated in a water bath for 20 min at 42°C. Then the absorbance was measured at 820 nm, and Pi content was calculated by comparison to standard curve. The Pi concentration was normalized to the shoot fresh weight.

### RNA isolation and qRT-PCR

RNA was isolated using the Qiagen plant RNA mini kit according to the manufacturer’s instructions, including an on column DNAse treatment. cDNA was made using the iScript cDNA Synthesis kit (Bio-Rad) using 300 ng total RNA as template. qRT-PCR was performed using the iQ SYBR Green Supermix (Bio-Rad) in a Bio-Rad CFX connect real-time system. Primers used for qPCR are listed in Table S1. Medicago Elongation factor 1 (*EF1*) was used as reference for normalization. Relative expression levels were calculated as 2^-^△△ct with three technical replicates for each sample.

### AM quantification

Mycorrhizal roots were stained using WGA-Alexafluor 488 (Thermo Fisher Scientific, USA). AM quantification was performed as described (McGONIGLE et al., 1990). In short, a Leica DM5500 B microscope equipped with an eyepiece crosshair was used to inspect the intersections between the crosshair and roots at 200x magnification. The following categories were noted in each intersection: root only, hyphopodium, extraradical hyphae, intracellular hyphae (Intrahyphae), good arbuscule (equal or larger than 40 μm), degrading arbuscule (less than 40 μm, and presence of septa). In cases where at one intersection more than one category was observed, then each category was counted once at that position. 100 intersections were inspected for each sample (containing 30 cm root) and the percentage of each category was calculated.

### Root exudate collection and quantification of strigolactones

Strigolactone analysis was done as described (van Zeijl et al., 2015; Liu et al., 2011). 6 seedlings of each genotype were grown in a X-stream 20 aeroponic system (Nutriculture) operating with 5 L of ½ Hoagland medium (Hoagland, 1950) containing 500 μM Pi in a greenhouse with natural light, 28 °C, 60% relative humidity. After 4 weeks, Pi starvation was initiated by replacing the high Pi medium with ½ Hoagland medium containing 20 μM Pi for one week. 24 hours before exudate collection, the medium was refreshed with new low Pi ½ Hoagland medium. The 5 liters containing 24-hour exudate were purified and concentrated by loading to a pre-equilibrated C18 column (Grace Pure C18-Fast 5000 mg/20 mL). Next, the column was washed with 50 mL of deionized water, followed by 50 mL of 30% acetone. Next, strigolactones were eluted with 50 mL 60% acetone and measured using a Quadcore as described (Kohlen et al., 2011).

### *R. irregularis* branching assay

*R. irregularis* spore germination and branching assays were performed as previously described (Besserer et al., 2006; Tsuzuki et al., 2016). 100 spores were grown on Solid M medium (BÉCARD and FORTIN, 1988) with 0.6% acetone, 0.01 μM GR24 or 100 times diluted root exudates (as collected above) in the dark at 22°C. 8 days after inoculation, hyphal branches were counted using a Leica M165 FC microscope.

### Confocal microscopy

The subcellular localization of fluorescently tagged proteins and detailed observation of the arbuscules using WGA-Alexafluor 488 stained roots were analysed using a Leica SP8 confocal microscope (For GFP/Alexafluor 488: excitation 488 nm, emission 500-540 nm; for DsRED/mCherry: excitation 552, emission 580-650 nm).

### Statistical analyses

For pairwise comparisons, data were analyzed using T-Test built in EXCEL with tail 1, type 2. One-way ANOVA built in Origin 2018 software was used to test difference over two groups of data with default settings. Replicates per experiment used are indicated in the corresponding figure legends.

## Acknowledgments

P.W. is supported by the China Scholarship Council (CSC) grant 201606310038 and by the Dutch research school Experimental Plant Sciences (EPS). We would like to thank Rene Geurts for critical reading and comments on the manuscript.

## Author Contributions

P.W. and E.L. conceived and designed experiments. P.W., R.S., and W.K.. performed experiments and/or data analyses. J.L. performed crossing. P.W., T.B. and E.L. wrote the manuscript.

## Competing Interest Statement

The authors declare no conflict of interest.

## Supplementary data

**Fig. S1.**
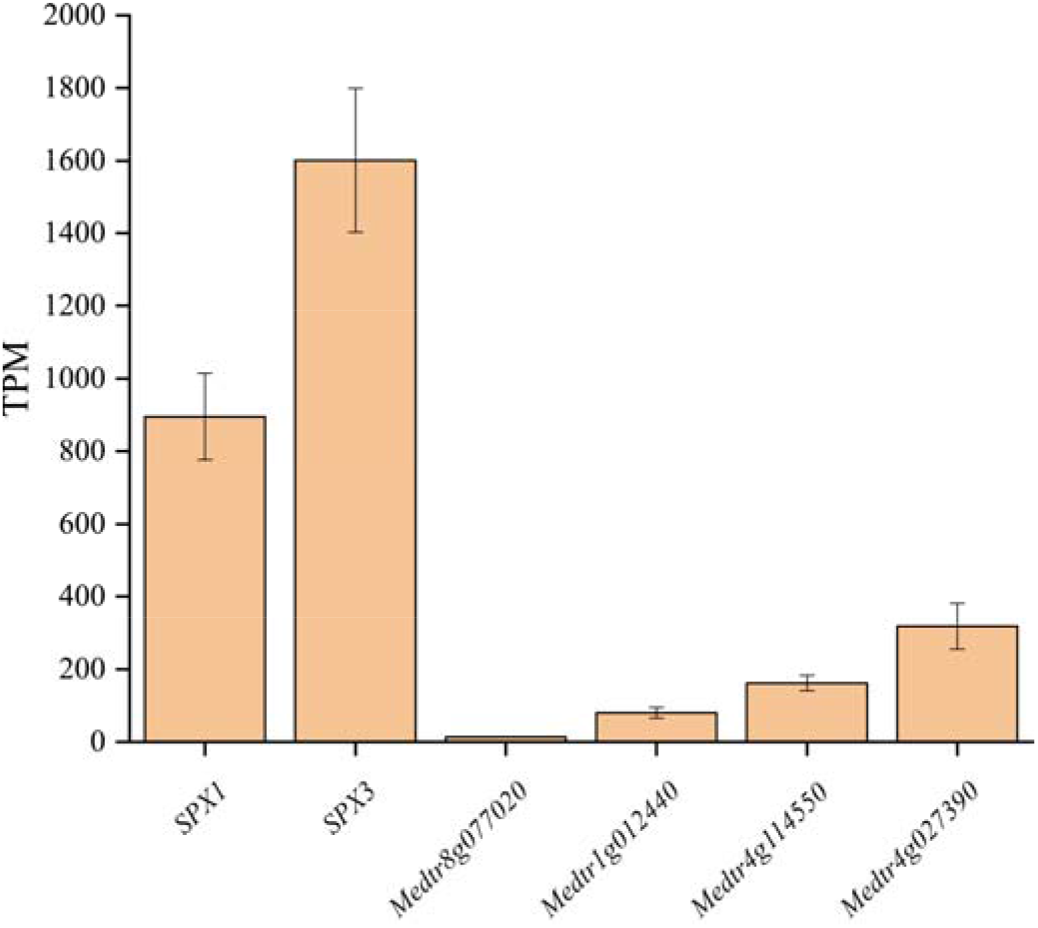
*SPX1* and *SPX3* are highly expressed in arbuscule-containing cells. RNAseq data of laser microdissected Medicago roots colonized by *R. irregularis* showing that *SPX1* and *SPX3* are the dominant SPX members expressed in arbuscule-containing cells. Data collected from (Zeng et al,. 2018). TPM= transcripts per million.

**Fig. S2.**
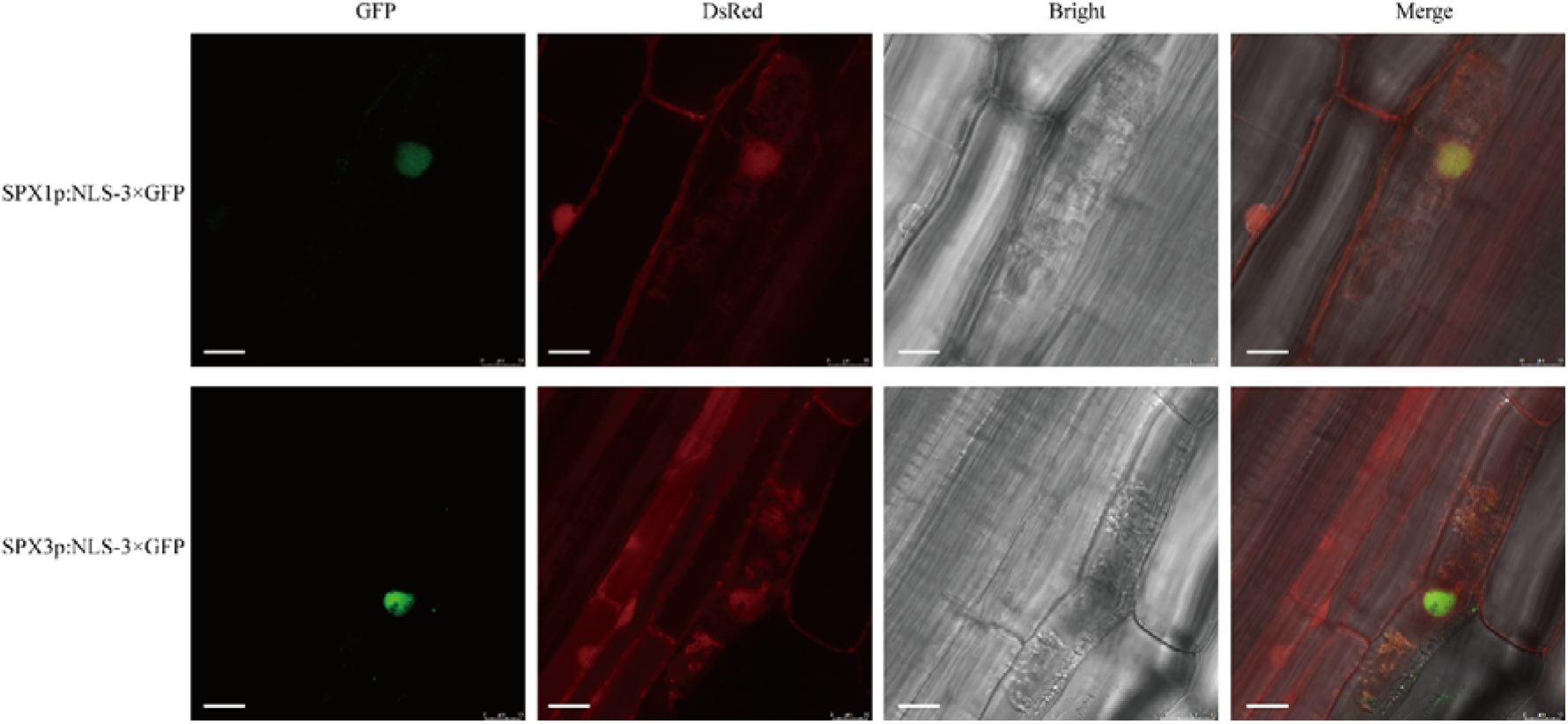
*SPX1* and *SPX3* specifically expressed in arbuscule-containing cell. Upper panels: Confocal images of nuclear localization signal (NLS)-3×GFP expressed from the *SPX1* promoter in mycorrhized *Medicago truncatula* roots. Bottom panels: Confocal images of NLS-3×GFP expressed from the *SPX3.* A co-expressed *UBp::DsRed* marker localizes to the nucleus and cytoplasm. Different panels represent the following channels (from left to right): GFP, DsRED, Bright field and all channels merged. GFP signal can only be detected in arbuscule-containing cell, while DsRed signal can be detected in neighboring non-colonized cortex cells. Scale bar = 10 μm.

**Fig. S3.**
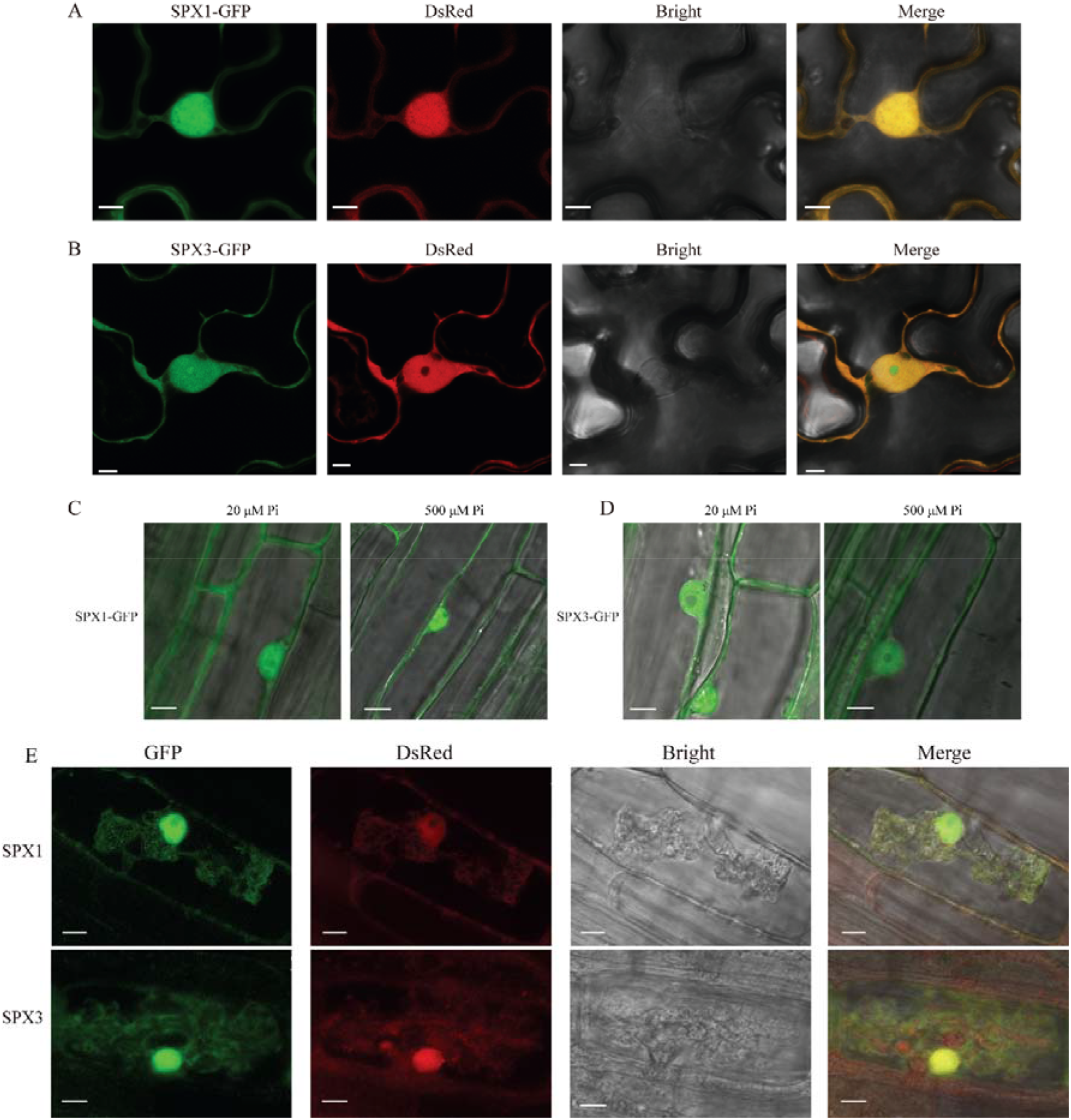
SPX1 and SPX3 localise to the nucleus and cytoplasm. (A) Confocal images of SPX1-GFP expressed from a constitutive *LjUbiquitin* promoter in *Nicotiana benthamiana* leaves. A co-expressed *UBp::DsRed* marker localizes to the nucleus and cytoplasm. Different panels represent the following channels (from left to right): GFP, DsRED, Bright field and all channels merged Scale bar = 8 μm. (B) Confocal images of *LjUBp::SPX3-GFP* in *Nicotiana benthamiana* leaves Scale bar = 5 μm. (C) Confocal images of *LjUBp::SPX1-GFP* in *Medicago truncatula* roots localizing to the nucleus and cytoplasm in low Pi and high Pi conditions. Scale bar = 10 μm. (D) Confocal images of *LjUBp::SPX3-GFP* expressed in *Medicago truncatula* roots in low Pi and high Pi conditions. Scale bar = 8 μm. (E) Confocal images of SPX1-GFP and SPX3-GFP expressed from their own promoter in *Medicago truncatula* roots in arbuscule-containing cells. Scale bar = 10 μm.

**Fig. S4.**
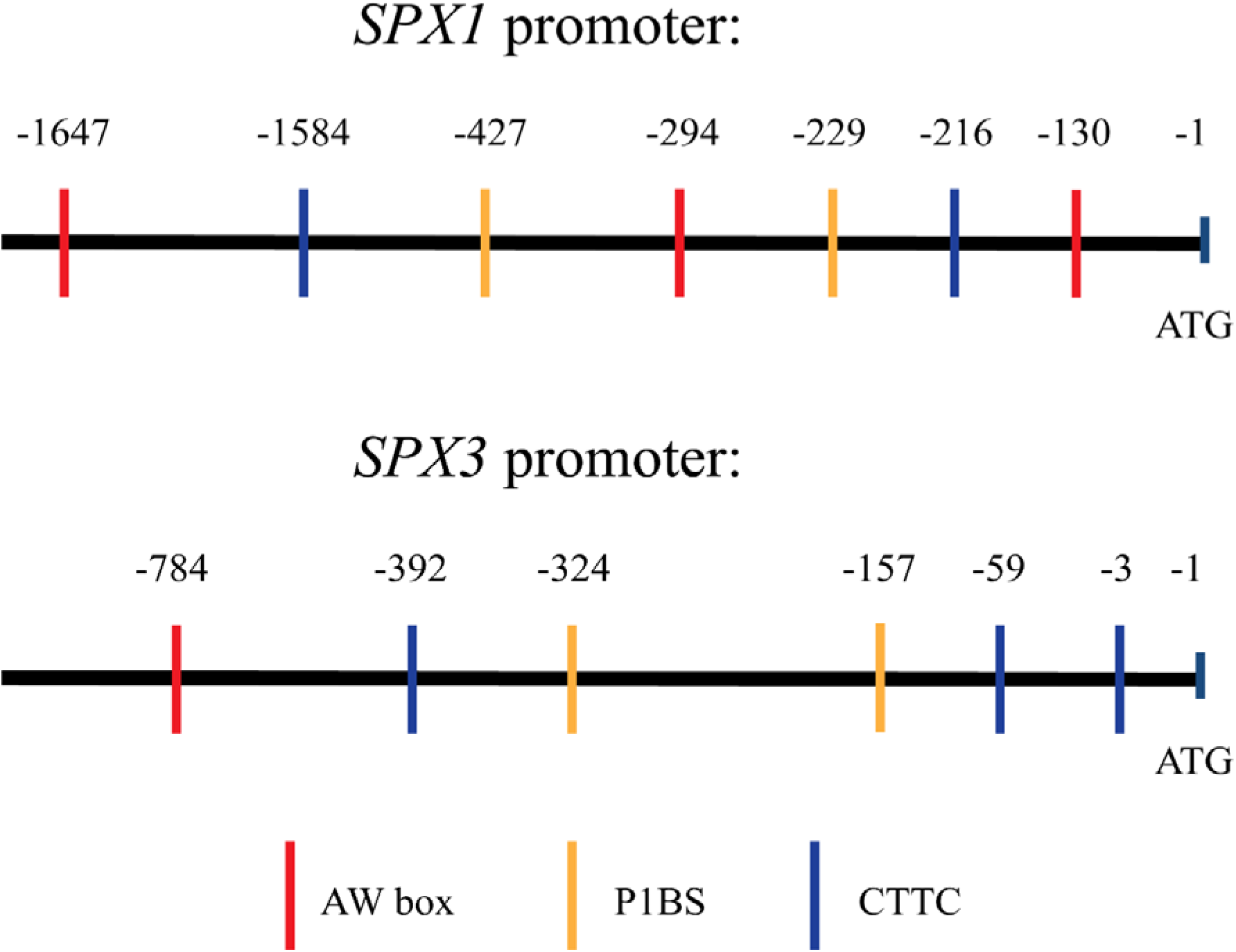
*SPX1* and *SPX3* promoters contain P1BS (GXATATXC), AW box (CG(X)7CXAXG) and CTTC (CTTCTTGTTC) *cis*-regulatory elements.

**Fig. S5.**
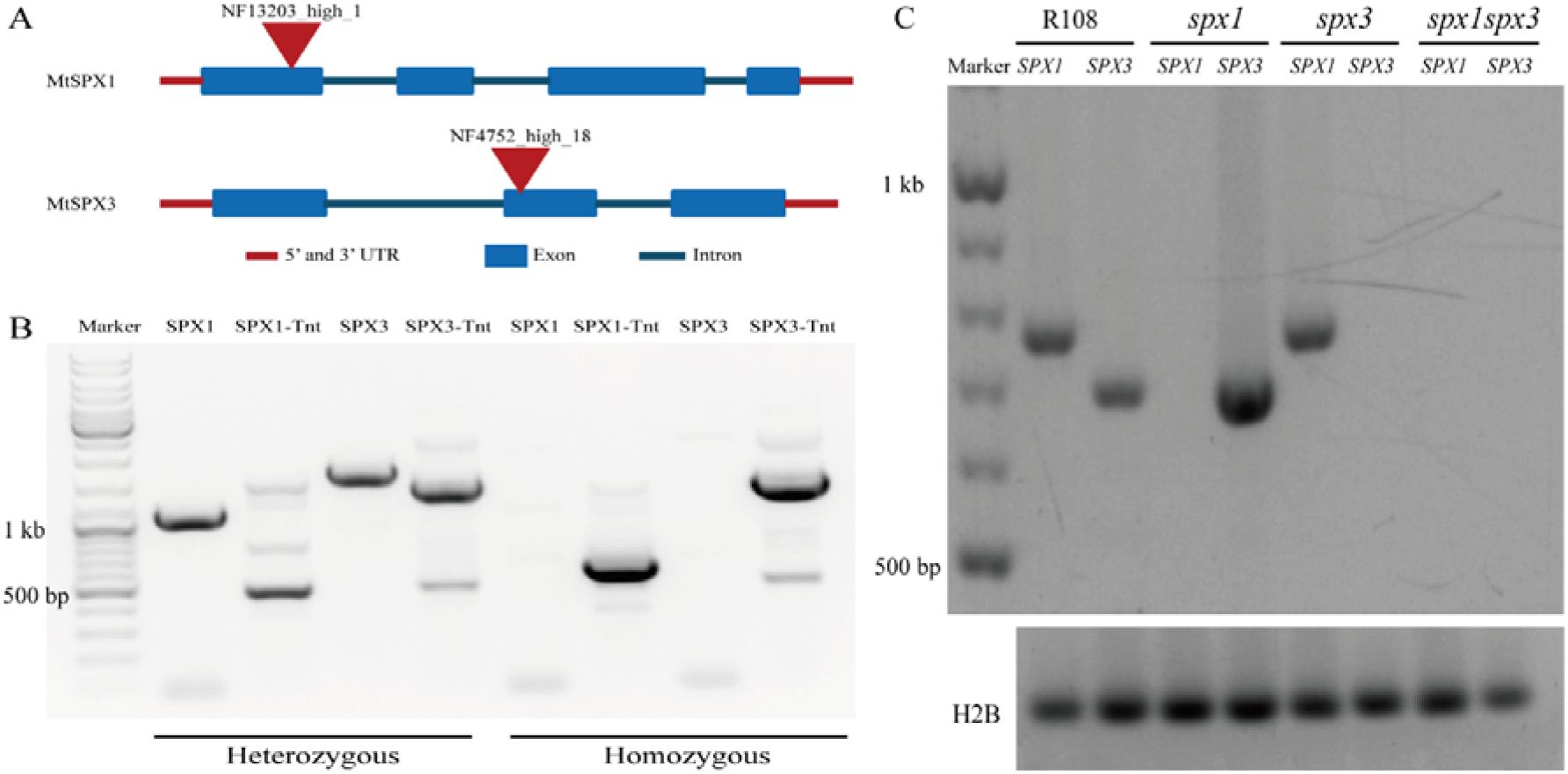
*spx1*, *spx3* and *spx1spx3* Tnt1-retrotransposon insertion lines. (A) Scheme of the Tnt1 insertions in *spx1* (NF13203) and *spx3* (NF4752), indicated by triangles. (B) PCR using genomic DNA as template confirming the Tnt1 insertions in both *SPX1* and *SPX3* in the *spx1spx3* double mutant. (C) RT-PCR showing the impairment of *SPX1* and *SPX3* expression in the respective mutants. *Histone 2B* (*H2B*) is used as control.

**Fig. S6.**
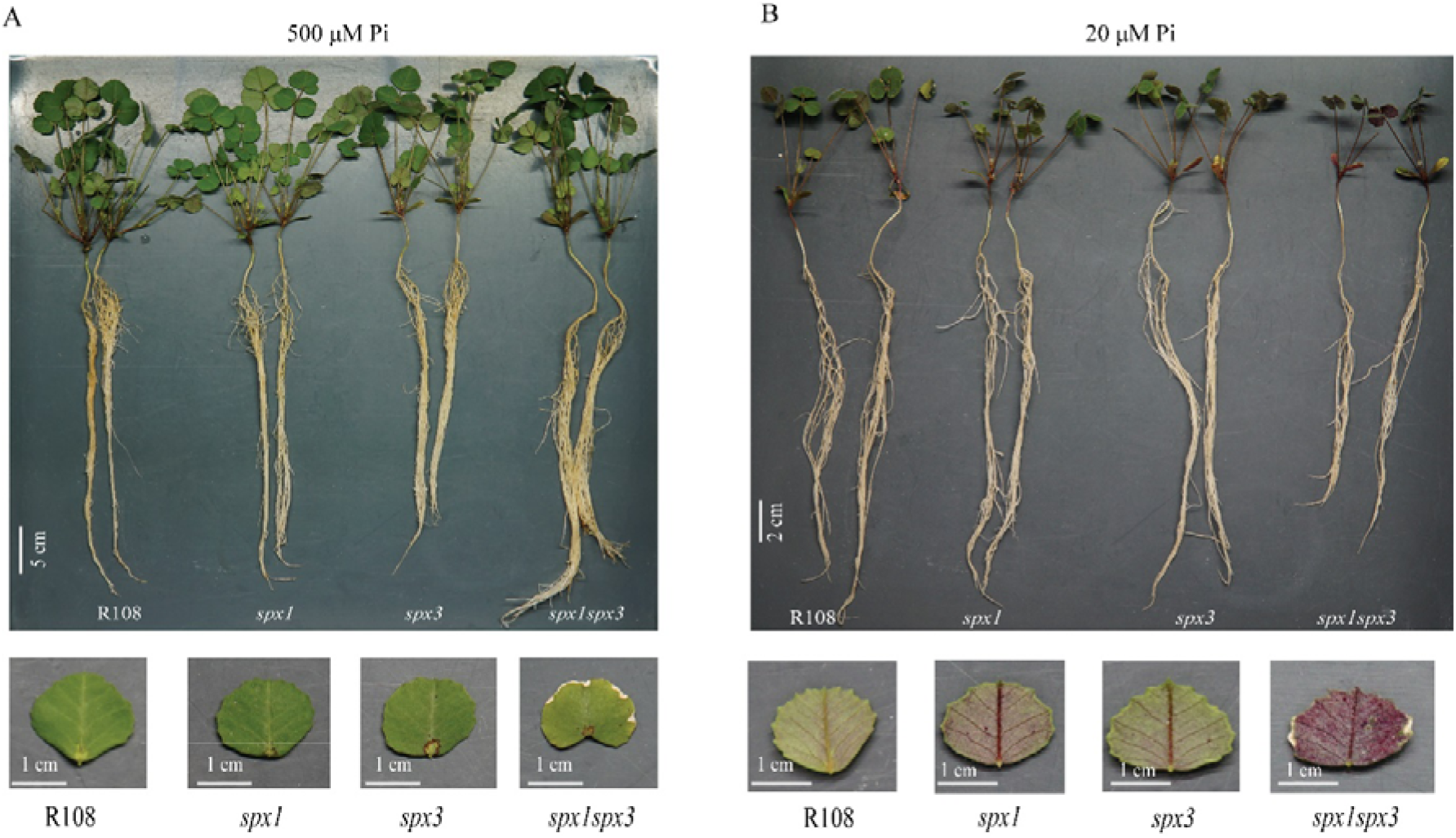
Phenotypes of *spx1*, *spx3*, and the double-mutant *spx1spx3*. (A) Phenotype of WT, *spx1*, *spx3*, and *spx1spx3* plants grown for 3 weeks in high Pi conditions (upper image). Bottom images show enlarged views of leaves. Note chlorosis of the leaf margins in the spx1spx3 double mutant. (B) Phenotype of WT, *spx1*, *spx3*, and *spx1spx3* plants grown for 3 weeks in low Pi conditions (upper image). The enlarged views in the bottom images show the accumulation of anthocyanins as purple coloration.

**Fig. S7.**
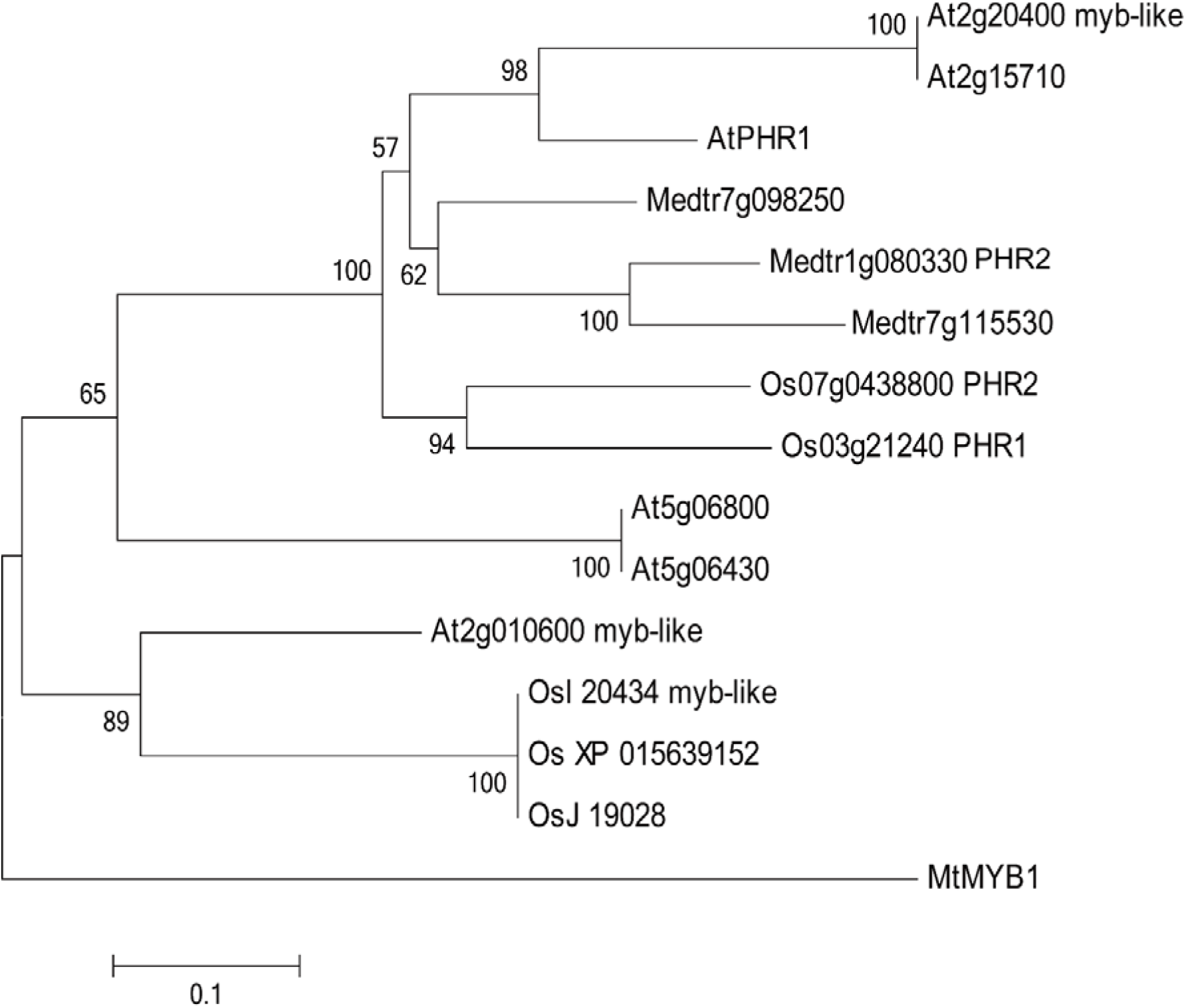
Phylogenetic tree of MYB family proteins from Medicago, Arabidopsis and rice generated using the neighbour-joining tree builder in Geneious R11.0 (https://www.geneious.com). 1000 bootstraps were used. Identifiers for the PHR genes are listed in Table S4.

**Fig. S8.**
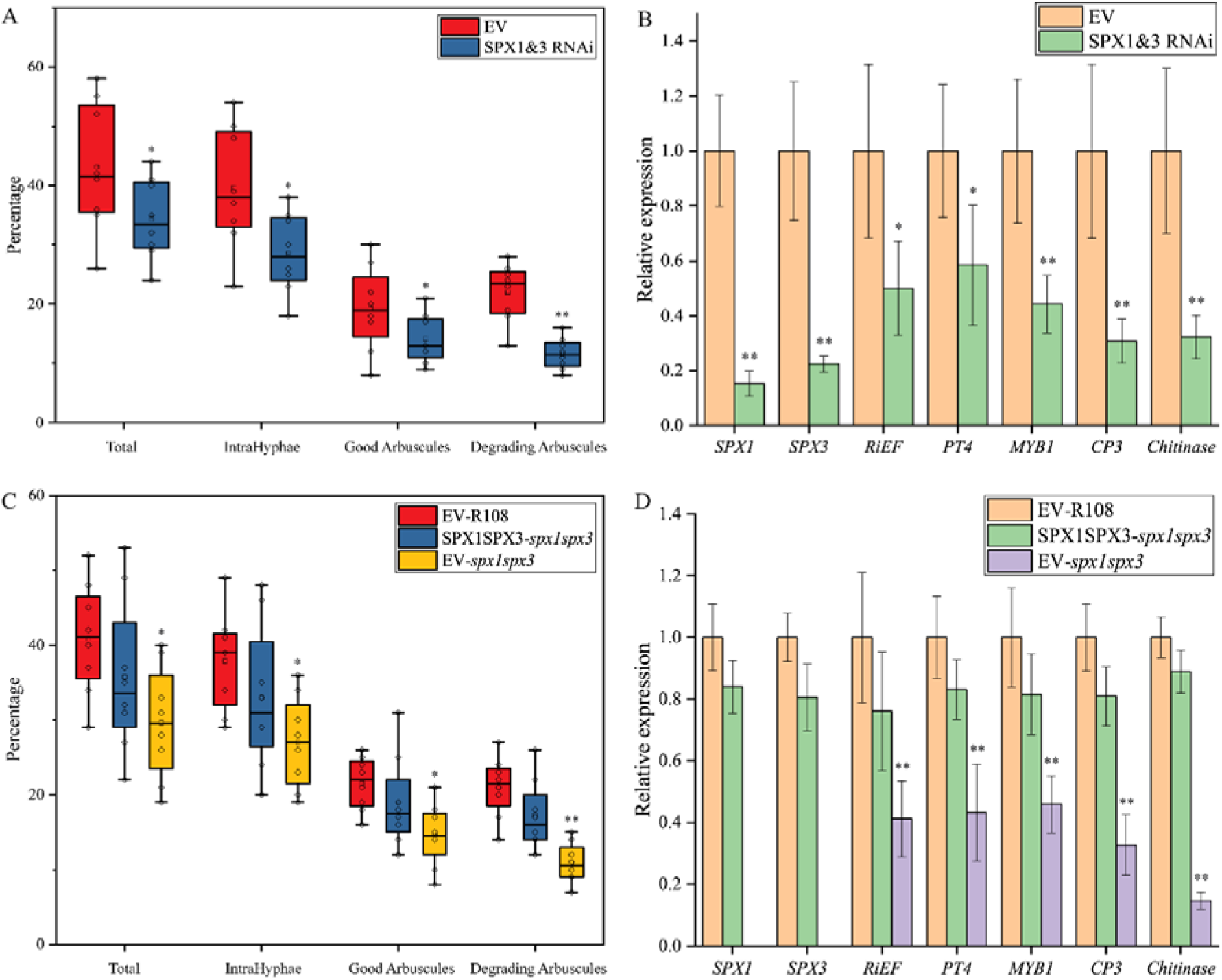
SPX1 and SPX3 control arbuscular mycorrhization. (A) RNAi knock down of *SPX1* and *SPX3* expression showing decreased arbuscular mycorrhization levels 3 weeks after inoculation in *Medicago truncatula* A17 roots compared to empty vector (EV) transformed roots. Significantly less degrading arbuscules were observed. 8 independent transformed roots were used as replicates for each sample. Total = presence of mycorrhiza in root segments, IntraHyphae = root segments containing intraradical hyphae; examples of good and degrading arbuscule classes is shown in Fig. 6 (B) Relative expression levels of *SPX1*, *SPX3*, *RiEF*, *PT4*, *CP3*, *Chitinase* and *MYB1* in root samples from (A) compared to empty vector (EV) controls. Values represent mean ± standard error of 6 independent replicates. (C) Expression of *SPX1* and *SPX3* under the control of their own promoters in the *spx1spx3* double mutant complemented the AM phenotype. 8 independent transformed plants were used as replicates for each sample. (D) Expression levels of *SPX1*, *SPX3*, *RiEF*, *PT4*, *CP3*, *Chitinase* and *MYB1* in root samples from (C). *MtEF1* was used as internal reference in the qPCR experiments shown in B and C. Values represent mean ± standard error of mean of 6 independent replicates. In (A-D) data significantly different from EV controls are indicated * P < 0.05; ** P < 0.01 (Student’s t-test).

**Fig. S9.**
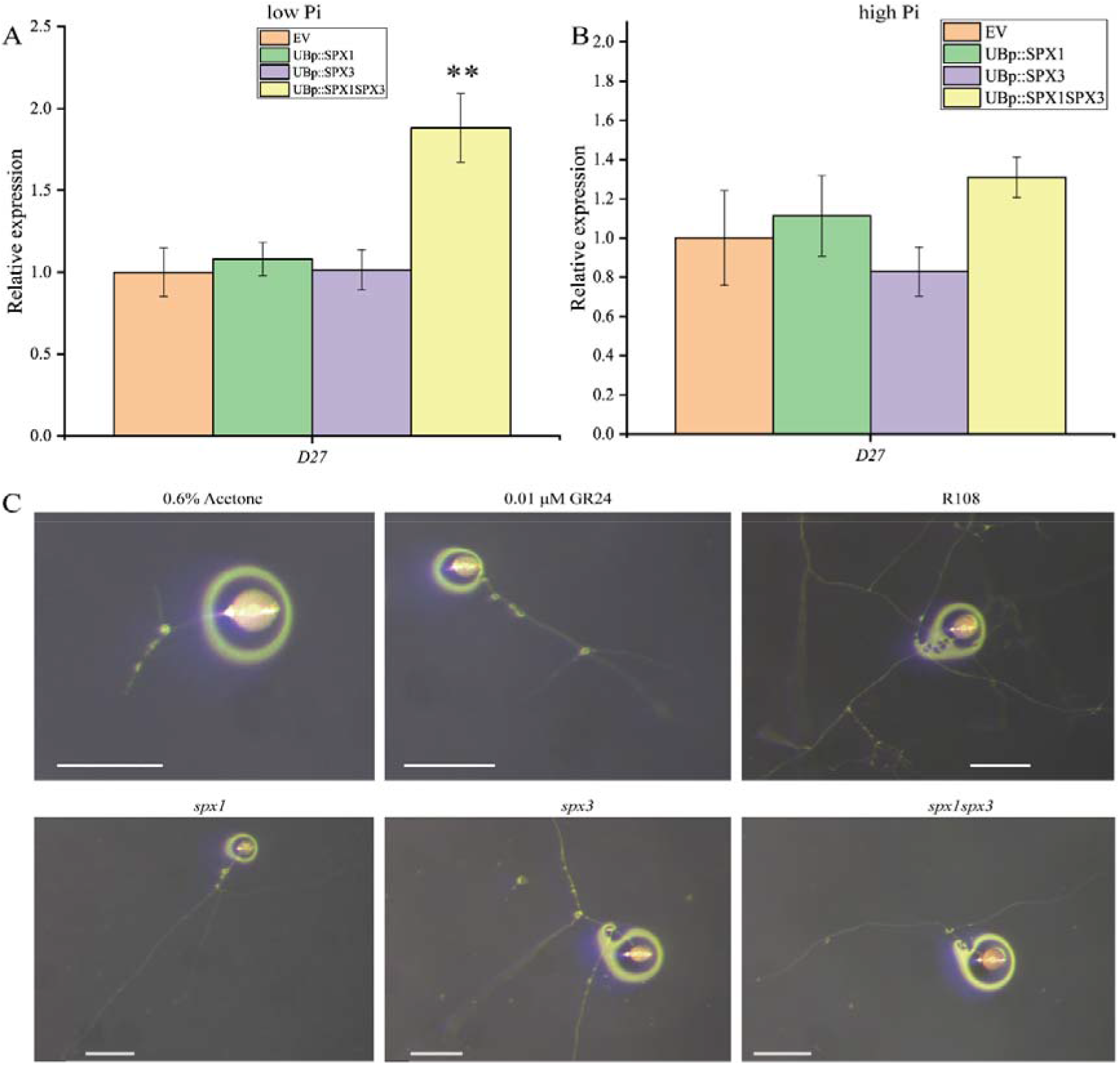
SPX1 and SPX3 likely regulate strigolactone biosynthesis. (A) Overexpression of *SPX1* and *SPX3* together in low Pi conditions induced *D27* expression. The samples correspond to Fig. 3E. (B) Overexpression of *SPX1* and *SPX3* in high Pi conditions did not affect *D27* expression. All values in (A-B) represent mean ± standard error of 3 independently transformed roots. *MtEF1* was used as reference gene in the qPCR experiment shown in A-B. Data significantly different from the corresponding empty vectors transformed controls are indicated ** P < 0.01 (Student’s t-test). (C) Representative images of *Rhizophagus irregularis* spores treated with 0.6% acetone, 0.01 μM GR24, roots exudates of R108, *spx1*, *spx3* and *spx1spx3*. Scale bar = 250 μm.

**Fig. S10.**
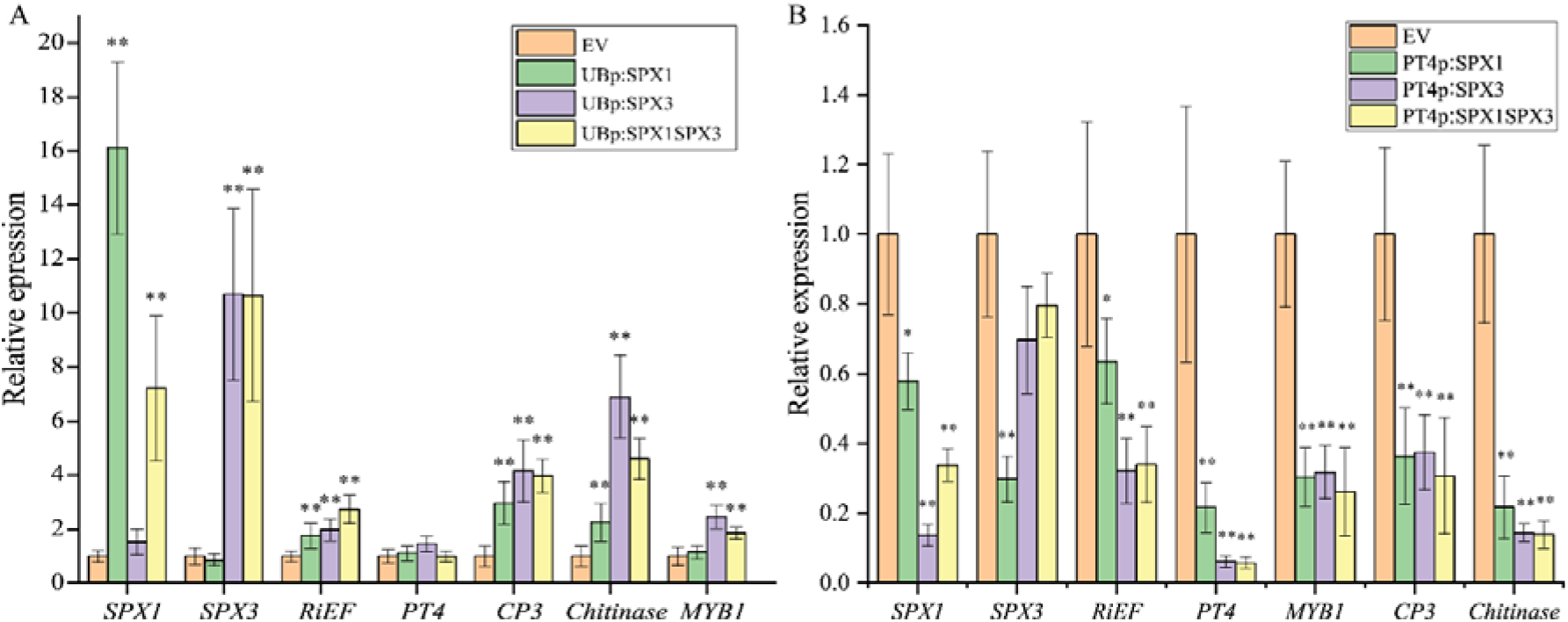
qPCR analysis of overexpression of *SPX1/3* under the control of the *LjUbiquitin1* or *PT4* promoter increased AM colonization and/or arbuscule degradation. (A) Expression levels of *SPX1*, *SPX3*, *RiEF*, *PT4*, *CP3*, *Chitinase*, *MYB1* in root samples from Fig. 6A as determined by qPCR. *MtEF1* was used as internal reference. (B) Expression levels of *SPX1*, *SPX3*, *RiEF*, *PT4*, *CP3*, *Chitinase*, *MYB1* in root samples from Fig. 6D as determined by qPCR. *MtEF1* was used as internal reference. All values in (A, B) represent mean ± standard error of mean of 6 independently transformed roots. Expression data significantly different from EV controls are indicated ** P < 0.01 (Student’s t-test).

**Fig. S11.**
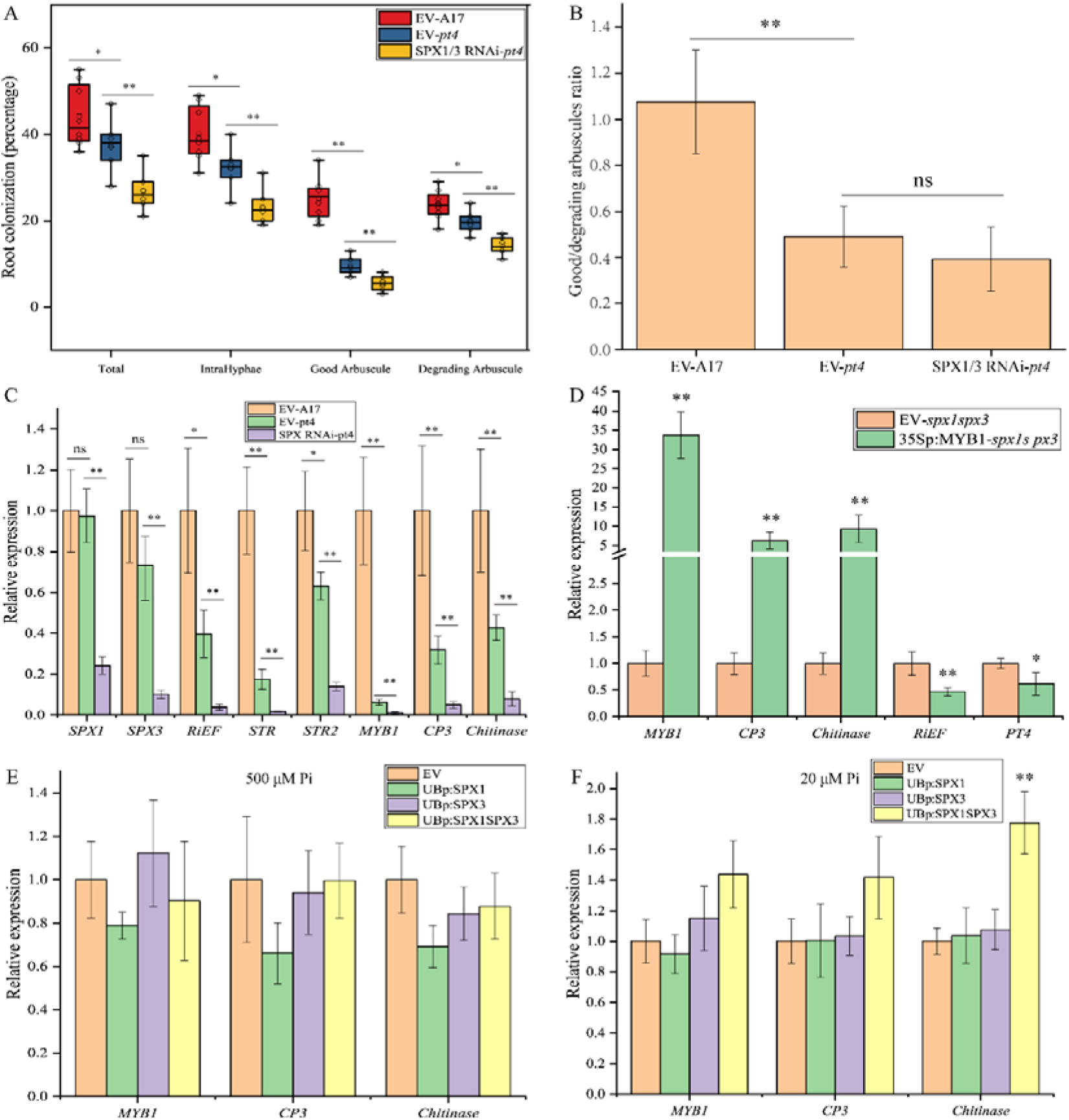
SPX1 and SPX3 function in relation to MYB1. (A) RNAi knock down of *SPX*1 and *SPX3* expression in the *pt4* mutant background decreased AM colonization but did not affect the good/degrading arbuscules ratio. 6 independently transformed plants were used as replicas for each sample. Data significantly different from indicated groups are indicated * P < 0.05; ** P < 0.01 (Student’s t-test). (B) Good-to-Degrading arbuscule ratio of mycorrhizal samples from (B). Values represent mean ± standard deviation of 6 independent replicates. Data significantly different from indicated groups are indicated ** P < 0.01; ns: not significant (Student’s t-test). (C) Relative expression levels of *SPX1*, *SPX3*, *RiEF*, *STR, STR2*, *MYB1*, *CP3* and *Chitinase* in samples from (A) compared to EV controls. Values represent mean ± standard error of 6 independently transformed roots. Data significantly different from indicated groups are indicated * P < 0.05; ** P < 0.01; ns: not significant (Student’s t-test). (D) Overexpression of *MYB1* under the control of *35S* promoter in the *spx1spx3* mutant background induced *CP3* and *Chitinase* expression and resulted in reduced *RiEF* and *PT4* expression compared to EV transformed roots 3 weeks post-inoculation. Values represent mean ± standard deviation of 3 independent replicates. Data significantly different from EV controls are indicated * P < 0.05; ** P < 0.01 (Student’s t-test). (E) Overexpression of *SPX1* and *SPX3* in high Pi conditions had no effect on *MYB1*, *CP3* and *Chitinase* expression. Values represent mean ± standard deviation of 3 independently transformed plants. (F) Overexpression of *SPX1* and *SPX3* together in low Pi conditions slightly increased *Chitinase* expression. cDNA samples used correspond to Fig. 2E. Values represent mean ± standard deviation of 3 independent replicates. *MtEF1* was used as internal reference in the qPCR experiments shown in C-F.

**Fig. S12.**
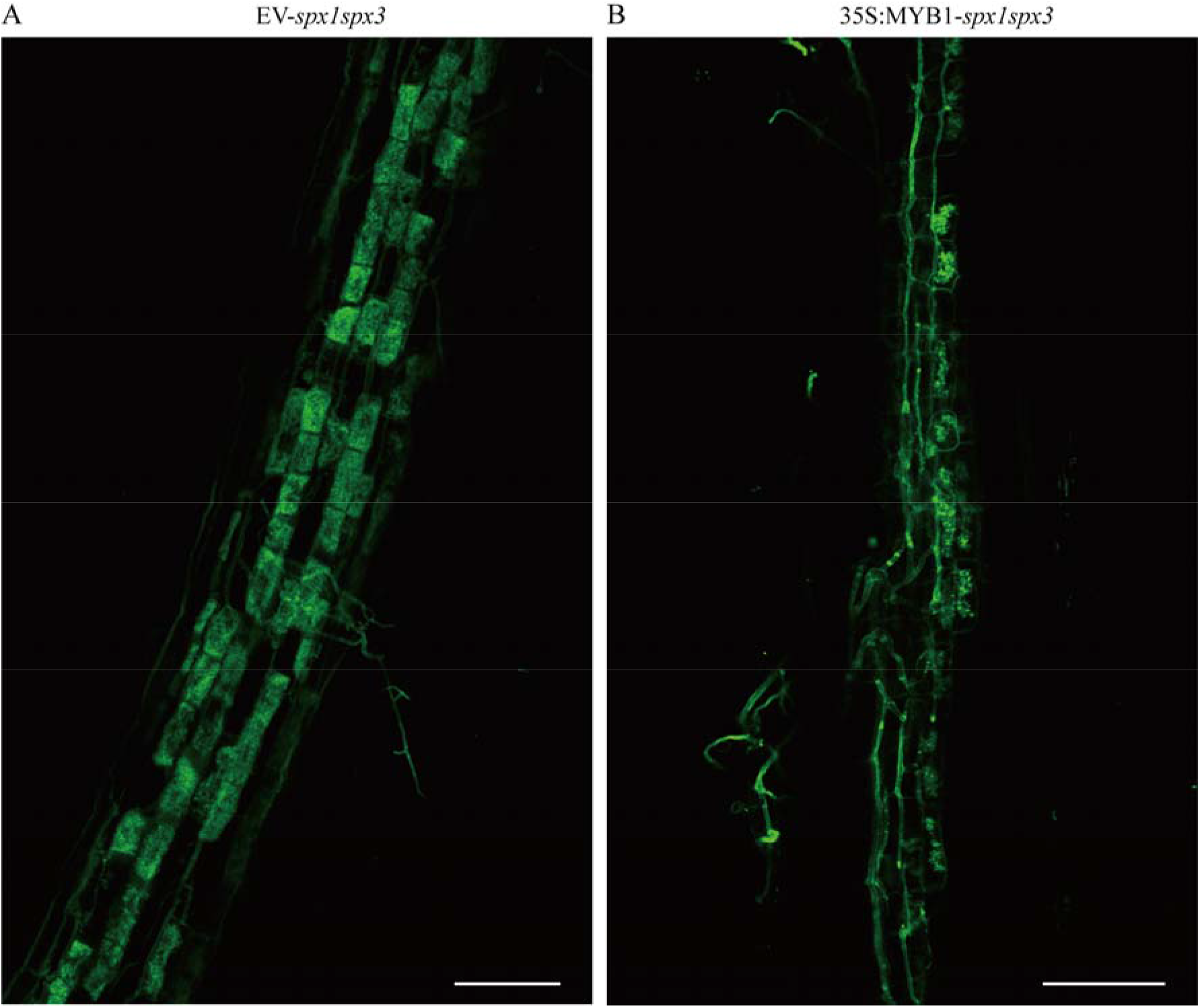
Representing images of *MYB1* overexpression mycorrhizal roots in the *spx1spx3* double mutant background. (A) Confocal picture of WGA-alexa488 stained empty vector transformed *spx1spx3* roots. Scale bar = 100 μm. (B) Confocal picture of WGA-alexa488 stained *35Sp:MYB1* transgenic *spx1spx3* roots. Scale bar = 100 μm.

## Supplementary Tables

**Table. S1.**
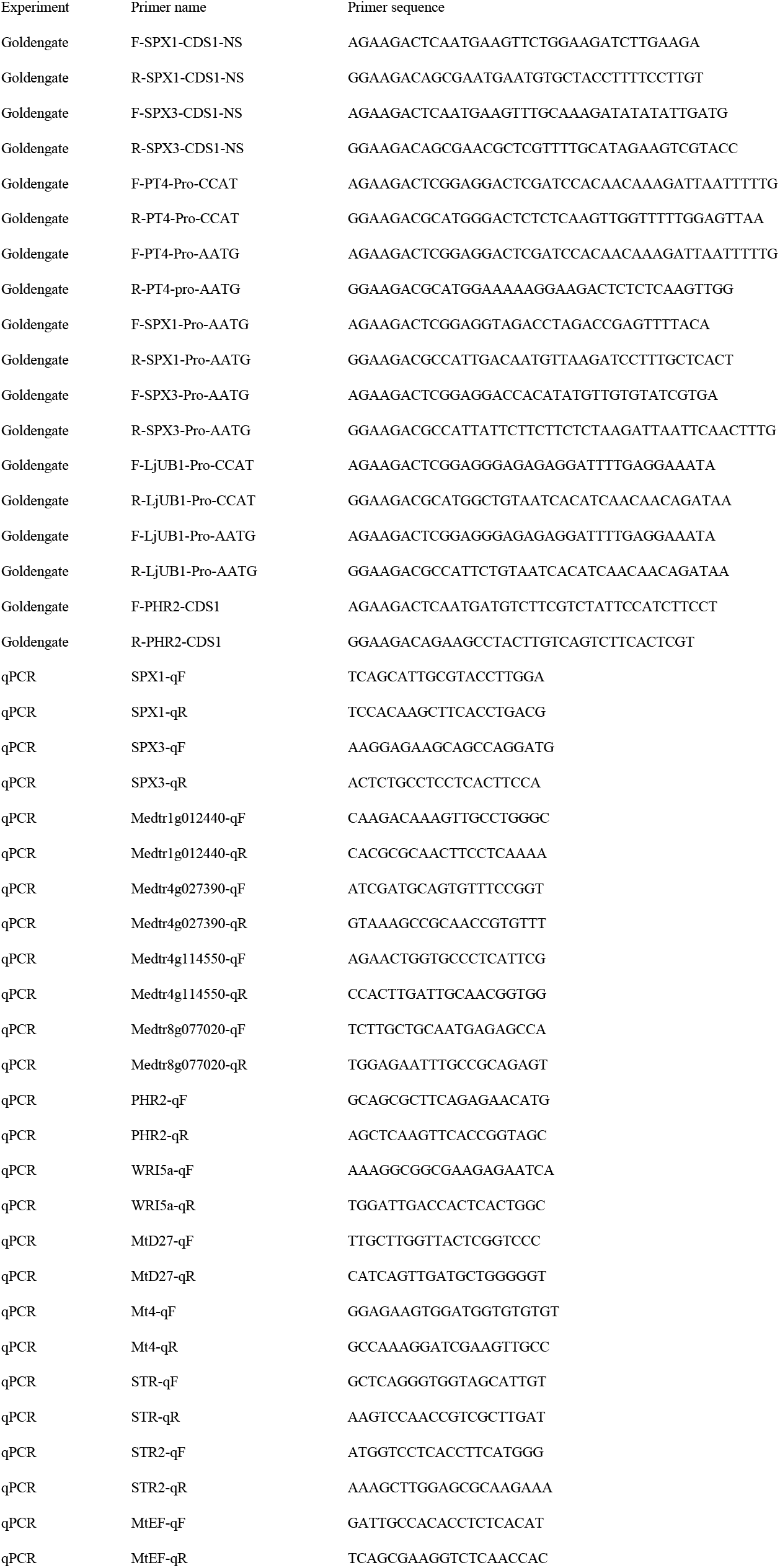
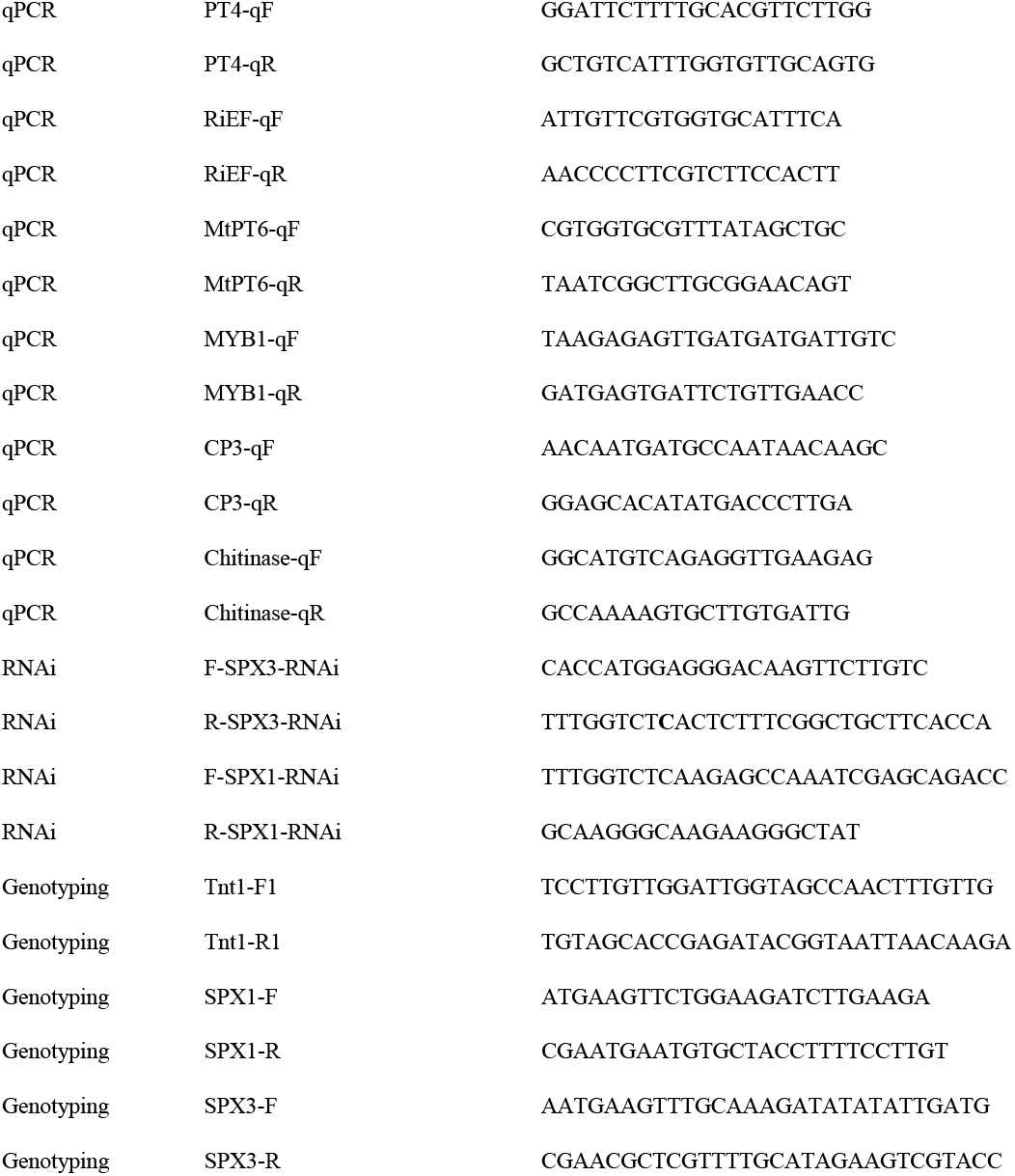
Primers list used in this study

**Table. S2.** Constructs made using the golden gate cloning system

| Level 0 | Level 1 | Level 2 |
| --- | --- | --- |
| pAGM1251-MtPT4p-CCAT | pICH47732-MtPT4p::SPX1-FLAG | pAGM4723-LjUBQ1p::SPX1-GFP-AtUBQ10p::DsRed |
| pICH41295-MtPT4p-AATG | pICH47751-MtPT4p::SPX3-FLAG | pAGM4723-LjUBQ1p::SPX3-GFP-AtUBQ10p::DsRed |
| pICH41295-LjUBQ1p-AATG | pICH47732-MtPT4p::FLAG-PHR2 | pAGM4723-LjUBQ1p::SPX1-FLAG-AtUBQ10p::DsRed |
| pAGM1251-LjUBQ1p-CCAT | pICH47732-LjUBQ1p::SPX1-FLAG | pAGM4723-LjUBQ1p::SPX3-FLAG-AtUBQ10p::DsRed |
| pICH41295-SPX1p-AATG | pICH47732-LjUBQ1p::SPX3-FLAG | pAGM4723-SPX1p::SPX1-GFP-AtUBQ10p::DsRed |
| pICH41295-SPX3p-AATG | pICH47732-LjUBQ1p::SPX1-GFP | pAGM4723-SPX3p::SPX3-GFP-AtUBQ10p::DsRed |
|  |  | pAGM4723-LjUBQ1p::SPX1-FLAG-AtUBQ10p::DsRed-LjUBQ1p::SPX3-FLAG |
| pAGM1287-SPX1 CDS1 no stop | pICH47732-LjUBQ1p::SPX3-GFP | pAGM4723-PT4p::SPX1-FLAG-AtUBQ10p::DsRed-PT4p::SPX3-FLAG |
| pAGM1287-SPX3 CDS1 no stop | pICH47751-LjUBQ1p::SPX3-GFP | pAGM4723-PT4p::SPX1-FLAG-AtUBQ10p::DsRed |
| pICH41308-PHR2 CDS1 | pICH47761-LjUBQ1p::GFP-MYB1 | pAGM4723-PT4p::SPX3-FLAG-AtUBQ10p::DsRed |
| pICH41308-MYB1-CDS1 | pICH47751-LjUBQ1p::GFP-PHR2 | pAGM4723-PT4p::FLAG-PHR2-AtUBQ10p::DsRed |
| pICH41308-NSP1-CDS1 | pICH47732-SPX1p::SPX1-GFP | pAGM4723-LjUBQ1p::FLAG-PHR2-AtUBQ10p::DsRed |
| pICH41308-DELLAΔ18-CDS1 | pICH47751-SPX3p::SPX3-GFP | pAGM4723-LjUBQ1p::GFP-PHR2-AtUBQ10p::DsRed |
|  | pICH47811-AtUBQ10p::DsRed | pAGM4723-LjUBQ1p::SPX1-FLAG-AtUBQ10p::DsRed-LjUBQ1p::GFP-PHR2 |
|  | pICH47751-LjUBQ1p::GFP-NSP1 | pAGM4723-LjUBQ1p::SPX3-FLAG-AtUBQ10p::DsRed-LjUBQ1p::GFP-PHR2 |
|  | pICH47732-LjUBQ1p::GFP-DELLAΔ18 | pAGM4723-LjUBQ1p::GFP-MYB1-AtUBQ10p::DsRed |
|  | pICH47751-LjUBQ1p::FLAG-NSP1 | pAGM4723-LjUBQ1p::GFP-PHR2-AtUBQ10p::DsRed |
|  |  | pAGM4723-LjUBQ1p::GFP-NSP1-AtUBQ10p::DsRed |
|  |  | pAGM4723-LjUBQ1p::SPX1-FLAG-AtUBQ10p::DsRed-LjUBQ1p::GFP-NSP1 |
|  |  | pAGM4723-LjUBQ1p::SPX3-FLAG-AtUBQ10p::DsRed-LjUBQ1p::GFP-NSP1 |
|  |  | pAGM4723-LjUBQ1p::GFP-DELLAΔ18-AtUBQ10p::DsRed |

**Table. S3.**
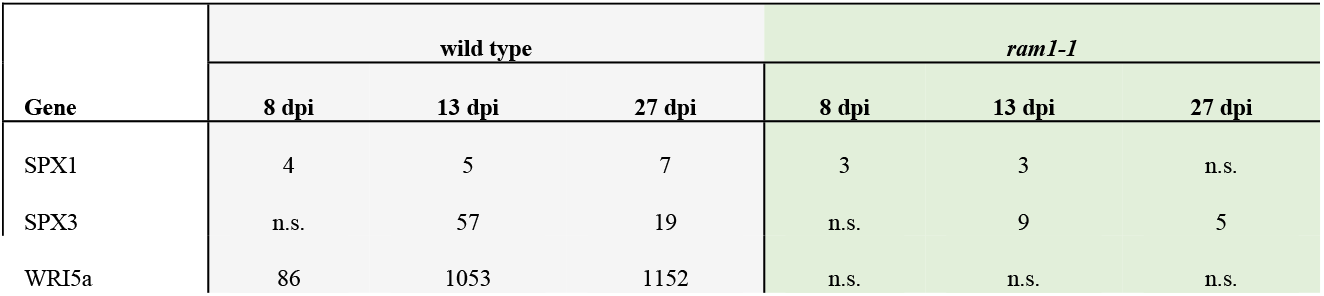
SPX1 and SPX3 expression fold changes in wild-type and *ram1-1* roots during mycorrhization. Fold changes with an FDR-corrected p-value < 0.05 are shown. Data collected from (Luginbuehl et al., 2017).

| Gene | wild type |  |  | *ram1-1* |  |  |
| --- | --- | --- | --- | --- | --- | --- |
|  | 8 dpi | 13 dpi | 27 dpi | 8 dpi | 13 dpi | 27 dpi |
| SPX1 | 4 | 5 | 7 | 3 | 3 | n.s. |
| SPX3 | n.s. | 57 | 19 | n.s. | 9 | 5 |
| WRI5a | 86 | 1053 | 1152 | n.s. | n.s. | n.s. |

**Table. S4.** Identifier of genes used for Phylogenetic tree

| Name | Gene ID |
| --- | --- |
| AtSPX1 | AT5G20150 |
| AtSPX2 | AT2G26660 |
| AtSPX3 | AT2G45130 |
| AtSPX4 | AT5G15330 |
| OsSPX1 | LOC_Os06g40120 |
| OsSPX2 | LOC_Os02g10780 |
| OsSPX3 | LOC_Os10g25310 |
| OsSPX4 | LOC_Os03g61200 |
| OsSPX5 | LOC_Os03g29250 |
| OsSPX6 | LOC9271158 |
| AtPHR1 | AT4G28610 |
| MtMYB1 | Medtr7g068600 |

## Footnotes

1 The author responsible for distribution of materials integral to the findings presented in this article in accordance with the policy described in the Instructions for Authors (www.plantcell.org) is: Erik Limpens.

